# Inhibition of p-glycoprotein does not increase the efficacy of proteasome inhibitors in multiple myeloma cells

**DOI:** 10.1101/2020.07.16.206102

**Authors:** Rachel L. Mynott, Craig T. Wallington-Beddoe

**Author notes:** **Corresponding author:** Dr Craig Wallington-Beddoe, Level 4, Flinders Centre for Innovation in Cancer Flinders University, Bedford Park, SA 5042, Australia.

## Abstract

The aim of this study is to determine whether manipulation of the drug transporter P-glycoprotein improves the efficacy of proteasome inhibitors in multiple myeloma cells. P-glycoprotein is a well-known drug transporter that is associated with chemotherapy resistance in a number of cancers but its role in modulating the efficacy of proteasome inhibitors in multiple myeloma is not well understood. Research has shown that the second generation proteasome inhibitor carfilzomib is a substrate of P-glycoprotein and as such its efficacy may correlate with P-glycoprotein activity. In contrast to carfilzomib, research concerning the first-in-class proteasome inhibitor bortezomib is inconsistent with some reports suggesting that inhibition of P-glycoprotein increases bortezomib cytotoxicity in multiple myeloma cells whereas others have shown no effect. Through the mining of publicly available gene expression microarrays of patient bone marrow, we show that P-glycoprotein gene expression increases with the disease stages leading to multiple myeloma. However, RNA-seq on LP-1 cells treated with bortezomib or carfilzomib demonstrated minimal basal P-glycoprotein expression which did not increase with treatment. Moreover, only one (KMS-18) of nine multiple myeloma cell lines expressed P-glycoprotein, including RPMI-8226 cells that are resistant to bortezomib or carfilzomib. We hypothesised that by inhibiting P-glycoprotein, multiple myeloma cell sensitivity to proteasome inhibitors would increase, thus providing a potential approach to improving responses and reversing resistance to these agents. However, the sensitivity of multiple myeloma cells lines to proteasome inhibition was not enhanced by inhibition of P-glycoprotein with the specific inhibitor tariquidar. In addition, targeting glucosylceramide synthase with eliglustat did not inhibit P-glycoprotein activity and also did not improve proteasome inhibitor efficacy except at a high concentration. We conclude that P-glycoprotein is poorly expressed in multiple myeloma cells, its inhibition does not enhance the efficacy of proteasome inhibitors, and it is unlikely to be a useful avenue for further translational research.

## Introduction

Multiple myeloma (MM) is an incurable malignancy of plasma cells (1). The malignant plasma cells usually produce monoclonal immunoglobulin and the disease is characterised by one or more of the four “CRAB” symptoms: high blood calcium, renal insufficiency, anaemia and bone lytic lesions (1). The median age at diagnosis is 69 years, meaning comorbidities and sub-optimal response to chemotherapies make the cancer difficult to treat in many patients (2). MM is currently incurable and if patients do not immediately present with chemotherapy resistance, they will eventually develop such resistance, contributing to a median overall survival of approximately 7 years (1).

P-glycoprotein (P-gp) is a well-known drug transporter that has been shown to play a role in drug resistance in a number of solid malignancies where it exports drugs, decreasing their intracellular concentration and thus reducing their efficacy (3, 4). Its relevance in chemotherapy resistance in MM was shown by Dalton *et. al.* where patients previously exposed to chemotherapy had a higher expression of P-gp on their plasma cells compared to newly-diagnosed patients (5). In the context of resistance to proteasome inhibitors, the most important class of novel therapies to treat MM, there have been conflicting reports in the literature. The second generation proteasome inhibitor carfilzomib is a known substrate of P-gp and its efficacy was shown to be enhanced in cancer cells, including MM, when P-gp is inhibited (6–9). In comparison, the efficacy of the first-in-class proteasome inhibitor bortezomib with P-gp inhibition was either improved (10–12) or not affected (6, 13–15) in MM and other human cancer cell lines. Improving the efficacy of proteasome inhibitors in MM and overcoming resistance to these agents would be very beneficial to improve treatment options for high-P-gp expressing patients.

The activity of P-gp can be inhibited directly, for example with the third-generation P-gp inhibitor tariquidar that blocks the ATPase activity of P-gp (8). However, P-gp can also be indirectly inhibited with research demonstrating that inhibition of the sphingolipid enzyme glucosylceramide synthase (GCS) reduces both gene and protein expression of P-gp (16, 17). GCS converts ceramide to glucosylceramide, the precursor of glycosphingolipids (GSLs) and it has been found that these GSLs upregulate P-gp gene expression via the *cSrc* and β-catenin signalling pathways in cancer cells (16). Eliglustat is a specific and potent inhibitor of GCS, with an EC_50_ in the low nanomolar range, and is used to treat Gaucher’s disease (18–20). As an inhibitor of GCS, it reduces glucosylceramide levels in cells and thus GSLs, causing the same reduction in P-gp expression as seen when GCS is either genetically or chemically inhibited by RNA interference or D,L-Threo-1-phenyl-2-palmitoylamino-3-morpholino-1-propanol (PPMP), respectively (21, 22). Therefore, eliglustat can be used to indirectly inhibit P-gp.

Here we report our negative findings, demonstrating P-gp does not appear to be expressed in most of a diverse range of MM cell lines, including one resistant to proteasome inhibitors, nor is its expression increased with proteasome inhibitor treatment. Moreover, inhibition of P-gp in MM cell lines does not enhance the efficacy of proteasome inhibitors and may therefore not be a useful avenue for further translational research. Overall, these *in vitro* cell line data support a lack of a role for P-pg in modulating the effects of proteasome inhibitors in MM.

## Materials and Methods

### Cell lines

RPMI-8226 WT (ATC CCL-155) human MM cell line was obtained from the American Type Culture Collection (Manassas, VA). Resistant RMPI-8226 V10R (bortezomib resistant) and RPMI-8226 C10R (carfilzomib resistant) cell lines were a gift from Professor Robert Orlowski (MD Anderson Cancer Centre, Houston, TX). LP-1 (ACC 41) human MM cell line was purchased from the Leibniz Institute DSMZ-German Collection of Microorganisms and Cell Cultures GmbH (Braunschweig, Germany). KMS-11 (JCRB1179) human MM cell line was purchased from Cell Bank Australia (Sydney, Australia). MM.1S, MM.1R, NCI-H929 were kindly provided by Prof. Andrew Spencer (Monash University, Vic, Australia) and KMS-18 cells kindly provided by Prof. Junia Melo (SA Pathology, Australia). Cells were cultured in RPMI 1640 (Gibco, Waltham, MA) supplemented with 10% foetal bovine serum, 50 units/mL penicillin, 0.25 mg/mL streptomycin, 2 mM L-glutamate, 1 mM sodium pyruvate and 15 mM Hepes buffer (all Gibco). Cells were incubated at 37°C in 5% CO_2_. All cell lines were genetically authenticated by the Australian Cancer Research Foundation (ACRF) Cancer Genomics Facility (Adelaide, Australia) and determined mycoplasma-free using the MycoProbe mycoplasma detection kit (CUL001B, R&D Systems, Minneapolis, MN).

### Drugs, reagents and antibodies

Bortezomib (product number 100100, lot number TZC60223), carfilzomib (200630, TZC50520) and eliglustat (329461, XPR70327) were purchased from MedKoo (Morrisville, NC). Tariquidar (S8028, S802801) was purchased from Selleck Chemicals (Houston, TX). Rhodamine 123 (Rho123, 83702), propidium iodide (PI) and cOmplete EDTA-free Protease Inhibitor Cocktail were purchased from Sigma Aldrich (St. Louis, MO). Tris UltraPure, hydrochloric acid, sodium chloride, glycerol, β-glycerophosphate and sodium fluoride were purchased from Chem Supply (Adelaide, Australia). FITC Annexin V apoptosis marker, Cat # 556419, AB_2665412 and 10X Annexin-V Binding Buffer were purchased from BD Biosciences (San Jose, CA). Precision Plus Protein All Blue and Kaleidoscope Pre-stained Protein Standards molecular weight ladders were purchased from Bio-Rad (Hercules, CA). Vectashield mounting media with 4′,6-diamidino-2-phenylindole (DAPI) was purchased from Vector Labs (Burlingame, CA).

FITC Mouse Anti-Human P-glycoprotein (CD243) monoclonal antibody (clone 17F9, Cat # 557002, AB_396548) and FITC Mouse IgG2b κ Isotype Control monoclonal antibody (clone 27-35, Cat # 555742, AB_396085) were purchased from BD Biosciences. MDR1/ABCB1 rabbit monoclonal antibody (clone E1Y7B, Cat # 13342, AB_2631176) was purchased from Cell Signaling (Beverly, MA). Anti-actin mouse monoclonal antibody (clone C4, Cat # MAB1501, AB_2223041) was purchased from Millipore (Burlington, MA). Anti-alpha tubulin mouse antibody loading control (clone DM1A, Cat # ab7291, AB_2241126) was purchased from Abcam (Cambridge, United Kingdom). Pierce Goat Anti-Rabbit IgG (H+L) peroxidase conjugated goat secondary antibody (Cat # 31460, AB_228341) and Pierce Goat Anti-Mouse IgG (H+L) peroxidase conjugated goat secondary antibody (Cat # 31430, AB_228307) were purchased from Invitrogen (Waltham, MA). Cy™5 AffiniPure Donkey Anti-Rabbit IgG (H+L), AB_2340607 and FITC AffiniPure Donkey Anti-Mouse IgG (H+L), AB_2335588 polyclonal secondary antibodies were purchased from Jackson Laboratories (Bar Harbor, ME).

### Cell viability by trypan blue

Trypan blue (1:2 dilution, Life Technologies, Waltham, MA) was mixed 1:1 with MM cell suspension. 10 μL of this mixture was pipetted onto a haemocytometer slide and cells within the central grid counted. Dead cells were observed as blue whereas live cells were unstained. The concentration of cells was determined by the following equation: *n* × *d* × 10,000 = cells/mL where *n* is the number of cells counted and *d* is the dilution factor (i.e. 2).

### Analysis of DNA microarrays

Publicly available gene expression datasets were used to assess P-gp gene expression in CD138-positive bone marrow plasma cells obtained from healthy donors, monoclonal gammopathy of undetermined significance (MGUS), smouldering MM (SMM), newly-diagnosed MM and relapsed MM patients. The following datasets were used: GSE13591 (normal, n=5, MGUS, n=11, MM, n=142) (23), E-MTAB-363 (normal, n=5, MGUS, n=5, MM, n=155) (24) and GSE6477 (normal, n=15, MGUS, n=22, smouldering MM, n=24, new MM, n=73, relapsed MM, n=28) (25). Excel (Microsoft Corporation, Redmond, WA) was used to compare differences between two groups using the Student t test (log_2_ normally distributed data for GSE13591 and E-MTAB-363 or untransformed data GSE6477).

### RNA sequencing and analysis

RNA extraction was undertaken using the RNeasy Mini Kit (QIAGEN, Hilden, Germany). Approximately 1 million cells (>80% viable) were treated with proteasome inhibitors for 6 and 16 hours before RNA was extracted following instructions provided by the kit. PolyA+ enriched RNAseq libraries were multiplexed and sequenced on the Illumina NextSeq 500 platform. Raw data, averaging 100 million reads per sample were analysed and quality checked using the FASTQC program (http://www.bioinformatics.babraham.ac.uk/projects/fastqc). Reads were mapped against the human (hg19) reference genomes for LP-1 cells using the STAR spliced alignment algorithm (26) (version2.7.2c with default parameters and -chimSeqmentMin 20, --quantMode GeneCounts) returning an average unique alignment rate of 80%. Differential expression analysis was evaluated from the TMM normalised gene counts using R (version 3.2.3) and edgeR (version 3.3) (27, 28) following protocols as described. Alignments were visualised and interrogated using the Integrative Genomics Viewer v2.3.804. Samples were processed and analysed the Australian Cancer Research Foundation Cancer Genomics Facility (Adelaide, Australia).

### Surface P-glycoprotein expression by flow cytometry

Approximately 100,000 – 500,000 MM cells were washed in 1X PBS to pellet. Cell pellets were stained with 2 μL FITC P-gp antibody or matched isotype for 20 minutes in the dark at room temperature. Cells were washed in PBS again to remove unbound antibody. Flow cytometric analysis was undertaken, and data collected from 10,000 events on the CytoFLEX S Flow Cytometer (Beckman Coulter, Brea, CA). CytExpert Software (Beckman Coulter) was used for data collection and analysis.

### Cell viability by flow cytometry

Approximately 100,000 – 500,000 MM cells were cultured with and without drugs of interest for up to 72 hours. Cell suspensions were washed in 1X phosphate buffered saline (PBS) and cell pellets stained with 2 μL FITC Annexin-V, 2 μL 1:50 PI and 96 μL 1X Annexin V Binding Buffer for 10 minutes in the dark at room temperature. Cell viability was determined by acquiring 10,000 events using a CytoFLEX S flow cytometer (Beckman Coulter) and analysis of cell viability performed using CytExpert Software (Beckman Coulter). Viable cells are those being dual Annexin-V FITC and PI negative.

### P-glycoprotein activity by Rhodamine 123 efflux

Approximately 100,000 – 500,000 MM cells were treated in tissue culture plates with proteasome inhibitors with and without tariquidar or eliglustat. After 22 hours, 200 ng/mL of the cell permeable P-gp substrate Rho123 was added to each well and the plates incubated at 37°C, 5% CO_2_ for the remaining 2 hours. Cells were then washed twice in 1X PBS and analysed by flow cytometry. Mean fluorescence intensity (MFI) was recorded from 10,000 events on the CytoFLEX S Flow Cytometer (Beckman Coulter). The Rho123 dye is excited by the blue laser (488 nm) and its emission is detected using a 525 nm filter. CytExpert Software (Beckman Coulter) was used for data collection and analysis and data were normalised to unstained, untreated controls.

### Western blotting

3 to 5 million cells were cultured with drugs of interest and then lysed in 10 mM Tris/HCl (pH 7.4), 137 mM NaCl containing 10% glycerol, 1% NP40, 10 mM β-glycerophosphate, 2 mM sodium fluoride and cOmplete EDTA-free protease inhibitor cocktail. This remained at 4°C for 10 minutes before centrifugation at 14,000 × g to clarify the lysate. Equal amounts of protein were loaded onto a Mini-PROTEAN TGX Precast Gel (Bio-Rad) and the Trans-Blot Turbo Transfer System was used to transfer protein onto a nitrocellulose membrane (Bio-Rad). Membranes were then blocked in 5% non-fat dairy milk powder in 50 mM Tris (pH 7.4), 154 mM NaCl and 1% Tween20 at room temperature for two hours. Primary and secondary antibodies were used as per the manufacturers’ instructions and membranes were imaged on the ChemiDoc MP (Bio-Rad) after incubation in Clarity Western ECL Substrate (Bio-Rad). MDR1/ABCB1 and anti-actin primary antibodies were used at a dilution of 1:1000 and 1:5000, respectively, and HRP-conjugated secondary antibodies were used at a dilution of 1:15,000. Image Lab Software (Bio-Rad) was used for densitometric analysis of Western blot images.

### Statistical Analysis

Comparisons between two groups were undertaken in Microsoft Excel using the Student t test with a p-value of less than 0.05 considered statistically significant. Synergistic effects of drug combinations were determined using the fractional product method (29) with a value of < −0.1 indicating synergy and > 0.1 antagonism.

## Results

### Changes in P-gp gene expression with MM disease progression and in response to proteasome inhibitors

Publicly available gene expression microarrays were used to analyse P-gp gene expression in CD138-positive purified bone marrow plasma cells. In the three gene expression microarray datasets analysed, newly-diagnosed and relapsed MM patients but not those patients with the precursor conditions monoclonal gammopathy of undetermined significance (MGUS) and smouldering MM (SMM) had higher P-gp gene expression when compared to healthy controls (Fig 1). Unlike previous findings (30), there was no statistically significant difference in P-gp gene expression when comparing newly-diagnosed MM patients to patients whose MM had relapsed (Fig 1C), however it appeared to be more highly expressed in MM patients compared to MGUS in two of the datasets (Fig 1A and C).

**Fig 1.**
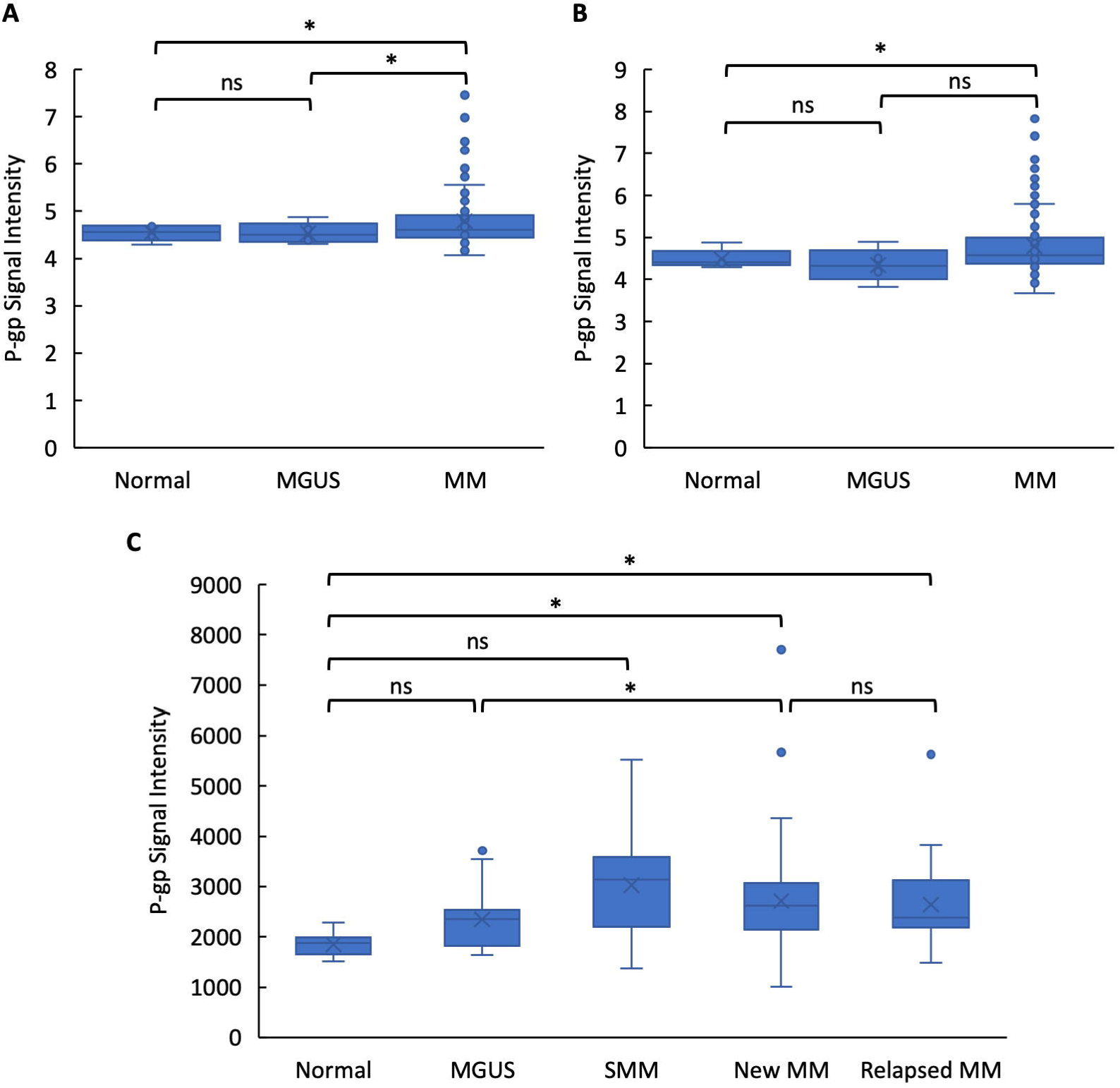
P-gp gene expression increases with disease progression in MM patients. Boxplots (median and interquartile range) showing P-gp expression from the GSE13591 (A), E-MTAB-363 (B) and GSE6477 (C) datasets. ns = not statistically significant, * p < 0.05. GSE13591 (normal, n=5, MGUS, n=11, MM, n=142), E-MTAB-363 (normal, n=5, MGUS, n=5, MM, n=155) and GSE6477 (normal, n=15, MGUS, n=22, smouldering MM, n=24, new MM, n=73, relapsed MM, n=28).

RNA-seq was undertaken to determine gene expression changes in LP-1 cells treated with 5 nM and 10 nM of bortezomib or carfilzomib for 6 hours. Analysis was undertaken for genes that had a count per million (CPM) of greater than 1 in at least two culture conditions. Unexpectedly, the data showed that P-gp was minimally expressed in this cell line (CPM < 1 for the control and proteasome inhibitor cultures) and so further normalisation and analysis was not undertaken for this gene.

### Basal P-gp protein expression in MM cell lines

Surface P-gp expression was analysed by flow cytometry. When compared to isotype, there appeared to be no P-gp expression on the surface of any of the human MM cell lines tested (Fig 2A). However, analysis of total cell P-gp by Western blot indicated that KMS-18 cells did express P-gp, whilst the other 8 MM cell lines examined, including the RPMI-8226 bortezomib (V10R) and carfilzomib (C10R) resistant cells, did not (Fig 2B). Due to the lack of P-gp in all MM cell lines aside from KMS-18, the latter cell line was used for most subsequent experiments exploring P-gp expression and activity.

**Fig 2.**
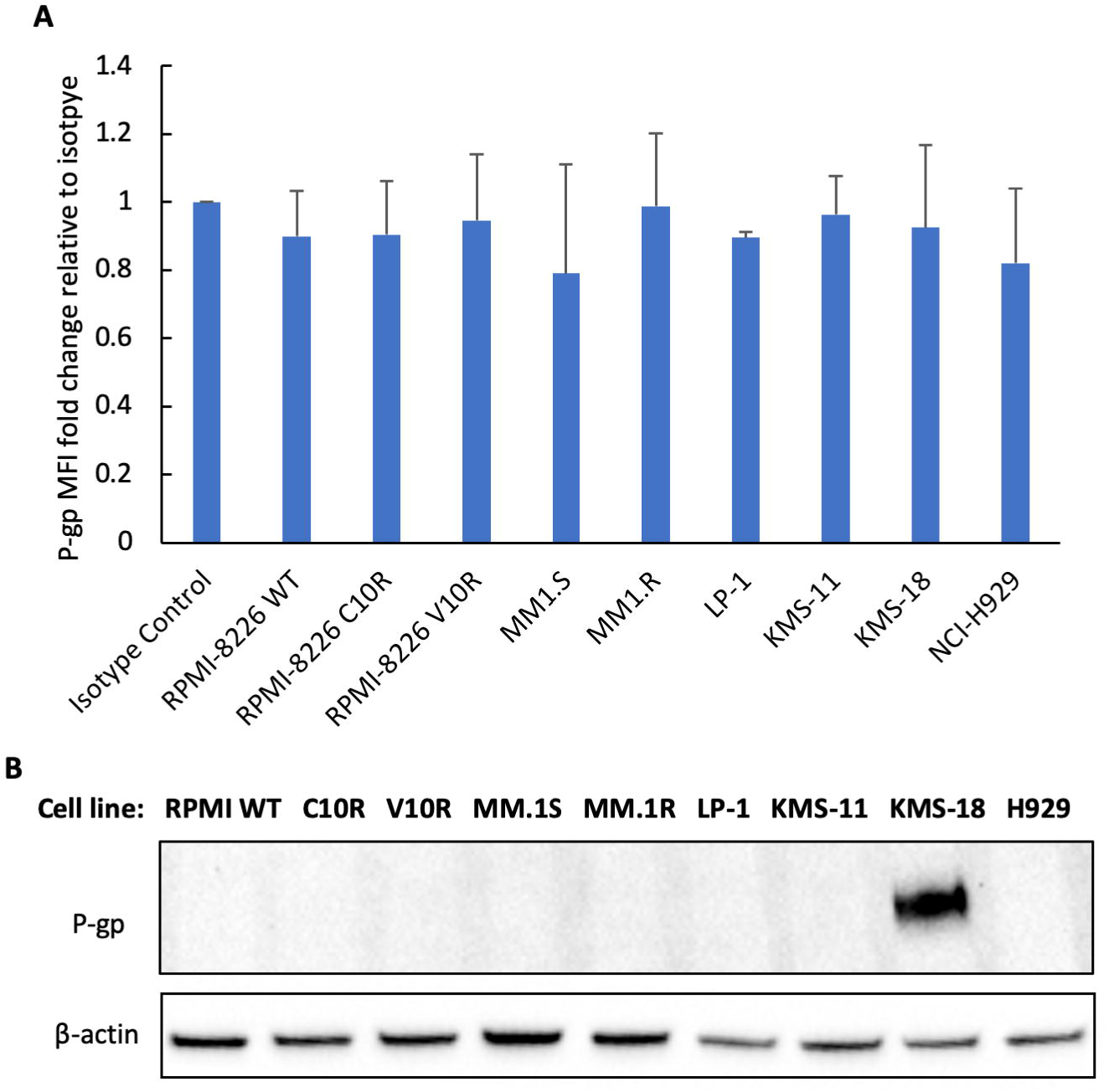
Basal P-gp in MM cell lines. A) Cell surface P-gp MFI of MM cell lines relative to isotype control. Data are mean ± standard deviation of duplicate measurements from three independent experiments. B) Western blot probed for total cell P-gp (170 kDa) in human MM cell lines. β-actin used as loading control. RPMI WT = RPMI-8226 WT, C10R = RPMI-8226 C10R, V10R = RPMI-8226 V10R and H929 = NCI-H929.

### Changes in P-gp expression, activity and MM cell viability in response to proteasome, P-gp and GCS inhibition

#### Proteasome inhibitors and changes in P-gp

LP-1 and KMS-18 MM cells cultured with bortezomib or carfilzomib for 24 hours did not appear to result in a clear increase in surface P-gp expression by flow cytometry (Fig 3A and C). Whilst both proteasome inhibitors demonstrated cytotoxicity, bortezomib appeared to have greater efficacy than carfilzomib in both LP-1 and KMS-18 cells despite carfilzomib being shown to be more effective than bortezomib in MM patients (Fig 3B and D) (31). Despite not causing an increase in surface P-gp in KMS-18 cells, bortezomib did result in total P-gp expression increasing after a 16-hour treatment (Fig 3E). Carfilzomib caused a more gradual increase in total P-gp expression, requiring a higher concentration to double the level of P-gp expression compared to the control culture (Fig 3F).

**Fig 3.**
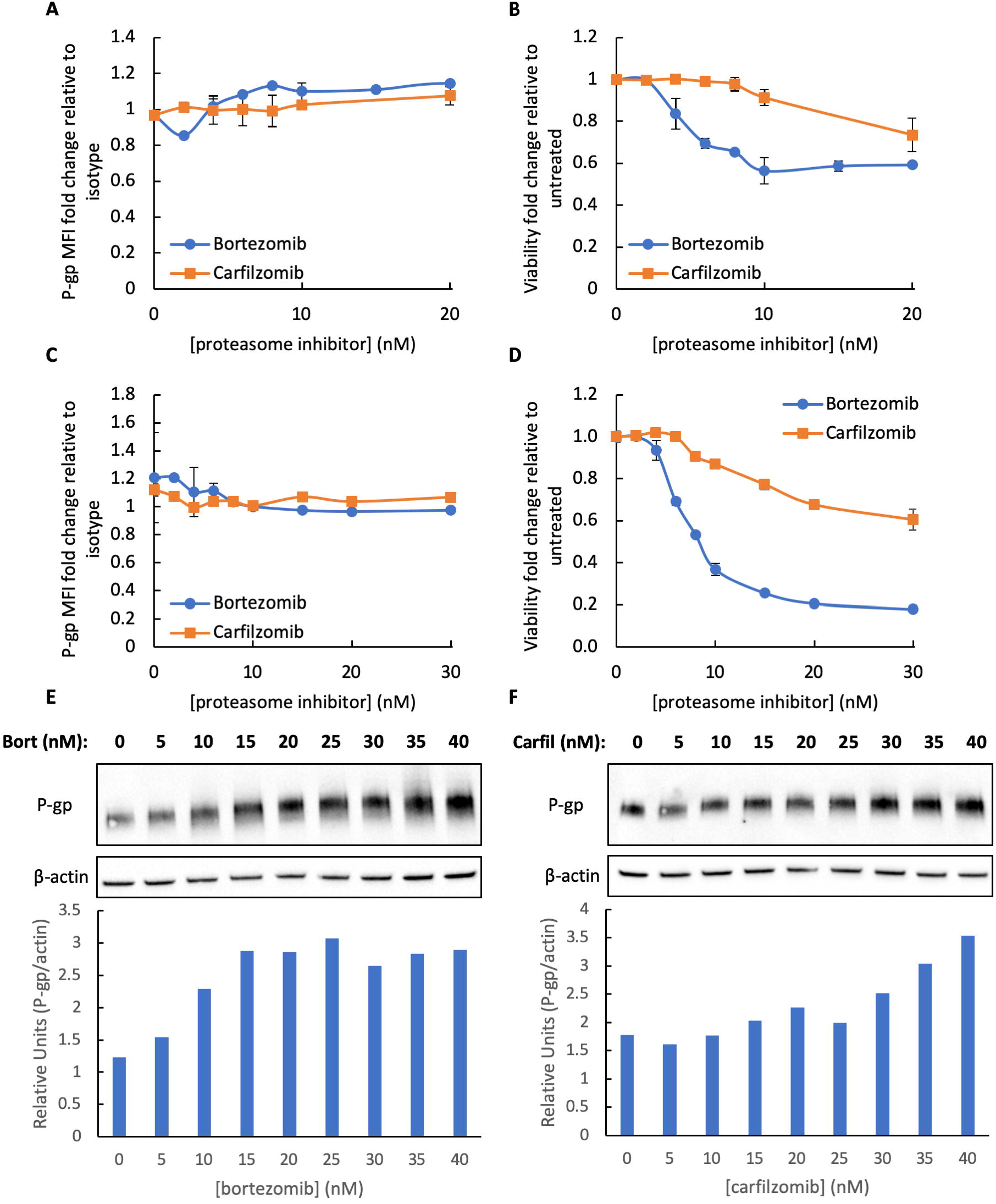
P-gp surface and total expression, and cell viability for MM cell lines treated with proteasome inhibitors. KMS-18 cells were treated with bortezomib or carfilzomib for 24 hours and flow cytometric analysis was undertaken to determine surface P-gp expression (A) and cell viability (B). LP-1 cells were treated with bortezomib or carfilzomib for 24 hours and flow cytometric analysis was undertaken to determine surface P-gp expression (C) and cell viability (D). Viability is represented as fold change relative to untreated control cultures. Data are mean ± standard deviation of duplicates from four biological replicates. KMS-18 cells were treated with bortezomib (E) or carfilzomib (F) for 16 hours and total P-gp expression was determined by Western blot. Associated densitometric analyses are displayed below each blot with P-gp band intensity relative to respective β-actin loading control.

In similar experiments examining proteasome inhibitor resistant MM cells, RPMI-8226 WT, RPMI-8226 V10R (bortezomib resistant) and RPMI-8226 C10R (carfilzomib resistant) cells, which all lack cell surface and whole cell P-gp (Fig 2A and B), showed no changes in surface P-gp expression after treatment with the proteasome inhibitors (Fig 4A and C). The expected sensitivity or resistance of each RPMI-8226 cell line to each proteasome inhibitor was confirmed (Fig 4B and D).

**Fig 4.**
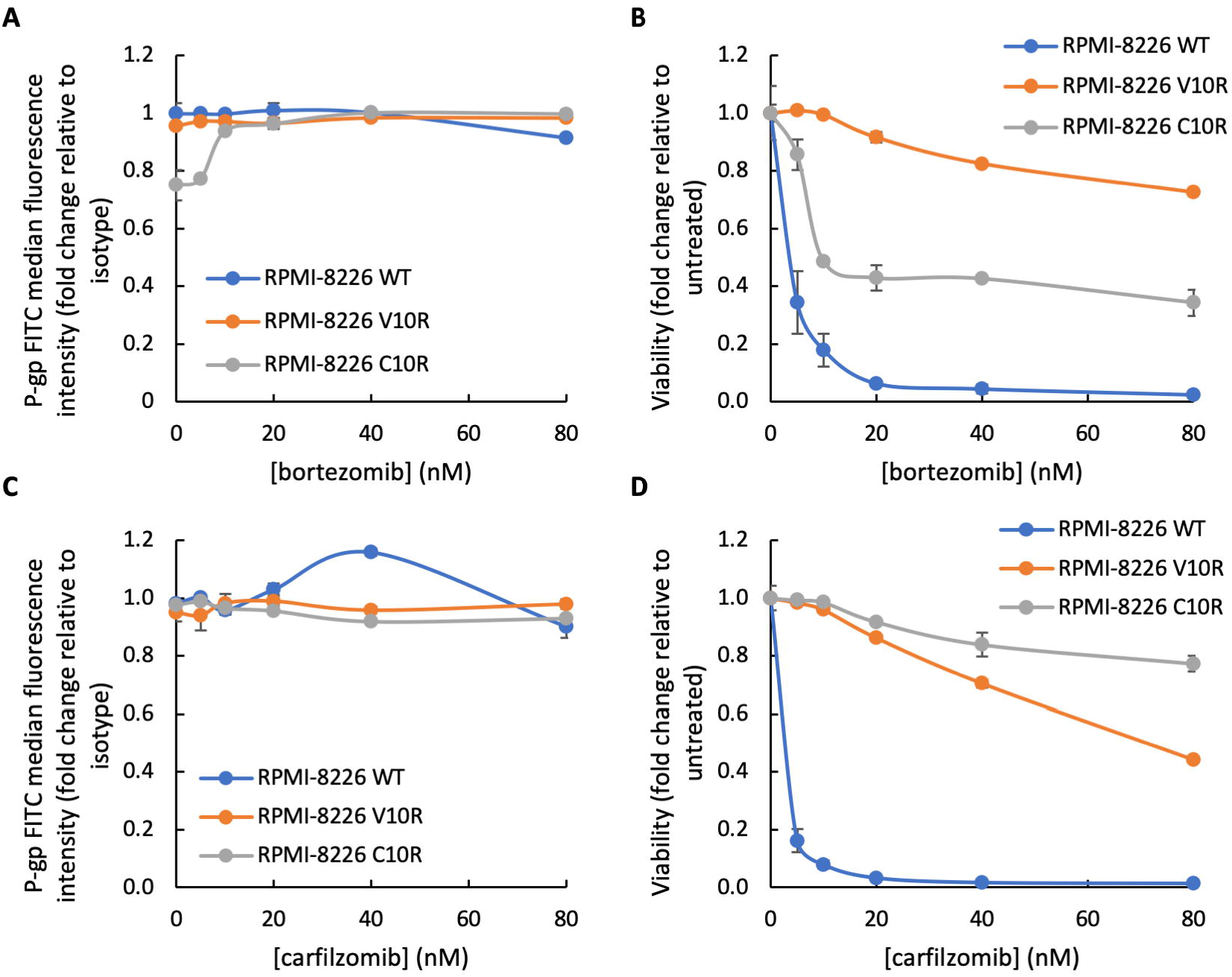
P-gp surface expression and cell viability of parental and proteasome inhibitor resistant RPMI-8226 MM cell lines treated with proteasome inhibitors. RPMI-8226 WT, V10R and C10R cells were treated with bortezomib or carfilzomib for 24 hours and flow cytometric analysis was undertaken to determine surface P-gp expression (A and C) and cell viability (B and D), respectively. Surface P-gp expression is represented relative to isotype and viability is represented as fold change relative to untreated control cultures. Data are mean ± standard deviation of duplicate measurements from three independent experiments.

### Inhibiting P-gp directly with tariquidar

P-gp protein expression changes in response to proteasome inhibitors with or without tariquidar were firstly assessed. Bortezomib alone caused an increase in total cell P-gp expression in KMS-18 cells which was not greatly altered by the addition of 1 μM tariquidar (Fig 5A and B), a concentration shown to effectively inhibit P-gp activity in cancer cells (32). Furthermore, the combination of bortezomib with tariquidar did not appear to cause a statistically significant change in surface P-gp expression compared to bortezomib alone (Fig 5C). Changes in P-gp activity in response to proteasome inhibitors with or without tariquidar were then examined. When KMS-18 cells were cultured with bortezomib alone, Rho123 MFI decreased with increasing concentrations of the drug, likely due to efflux of Rho123 from the cell. The addition of tariquidar inhibited the activity of P-gp as expected, causing an accumulation of Rho123 as seen by Rho123 MFI remaining constant with increasing concentrations of bortezomib (Fig 5D). Tariquidar did not appear to further sensitise cells to the cytotoxic effects of bortezomib, as neither an additive nor synergistic increase in cell death was seen in KMS-18 cells treated with the combination of the two drugs (Fig 5E).

**Fig 5.**
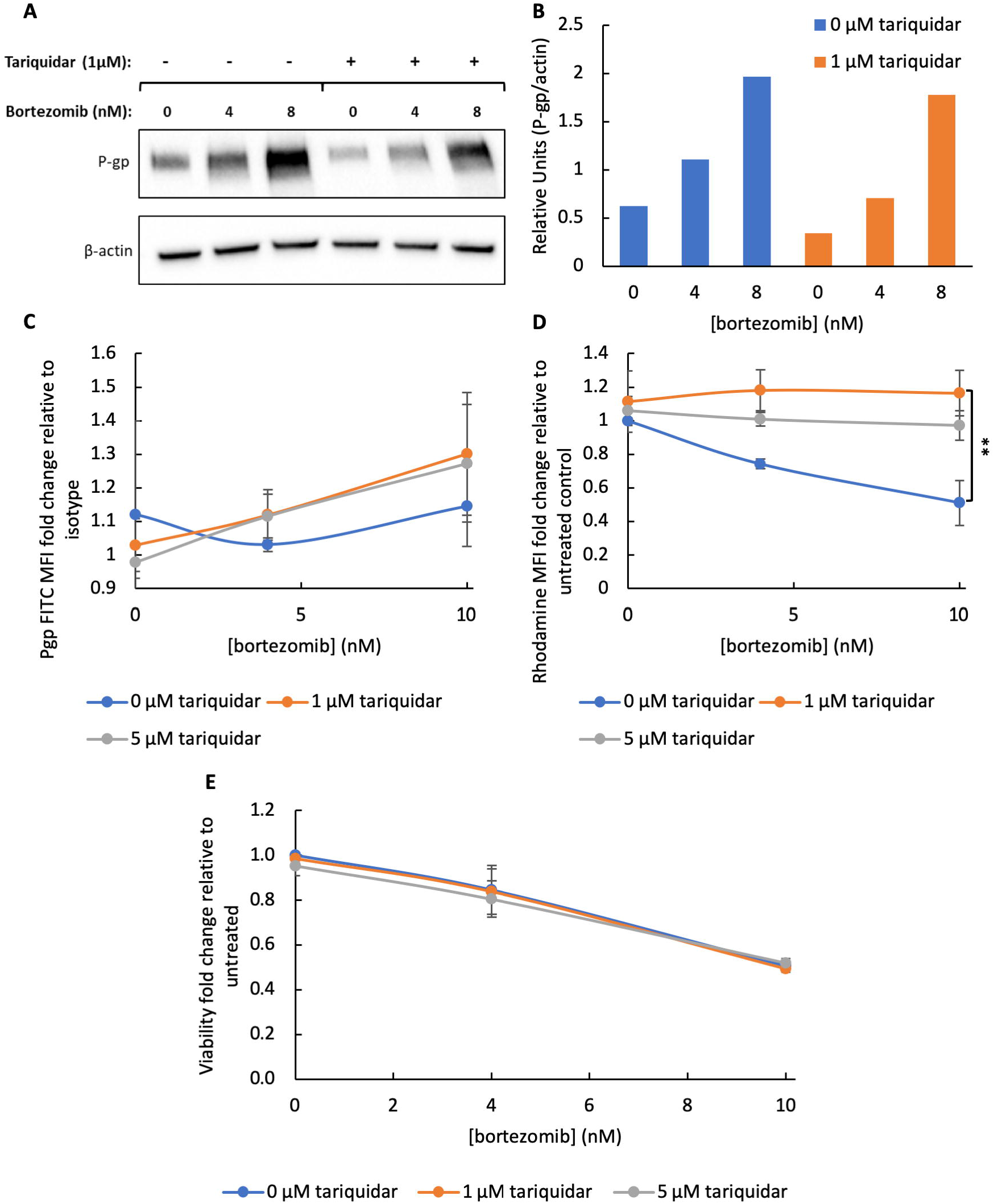
Tariquidar functionally inhibits P-gp but does not synergise with bortezomib. A) Western blot for KMS-18 cells treated with bortezomib and 0 or 1 μM tariquidar for 16 hours. B) Densitometric analysis of the Western blot displayed in (A), P-gp band intensity relative to respective β-actin loading control. KMS-18 cells were treated with bortezomib and 0, 1 or 5 μM tariquidar for 24 hours. Surface P-gp expression relative to isotype (C), Rho123 MFI normalised to unstained and relative to untreated (first data point of blue line) controls (D) and viability relative to untreated control (E) were analysed by flow cytometry. Flow cytometry data are mean ± standard deviation from three independent experiments. ** p < 0.01.

Carfilzomib alone did not appear to greatly change total P-gp expression in KMS-18 cells at the concentrations tested, however, the addition of 1 μM tariquidar caused a slight decrease (Fig 6A and B). On the cell surface, 5 μM tariquidar resulted in a small increase in P-gp expression when combined with 20 nM carfilzomib (Fig 6C). When KMS-18 cells were cultured with carfilzomib alone, the Rho123 MFI decreased with increasing concentrations of the drug, suggesting an increase in efflux of the dye. The treatments combining carfilzomib with tariquidar resulted in a much smaller decrease in Rho123 MFI, suggesting the addition of tariquidar causes an accumulation of dye in the cells when compared to carfilzomib alone, consistent with functional inhibition of P-gp by tariquidar (Fig 6D). As with bortezomib, the addition of tariquidar did not appear to further sensitise cells to the cytotoxic effects of carfilzomib, as neither an additive nor synergistic reduction in cell viability was observed in cells treated with the combination of the two drugs (Fig 6E).

**Fig 6.**
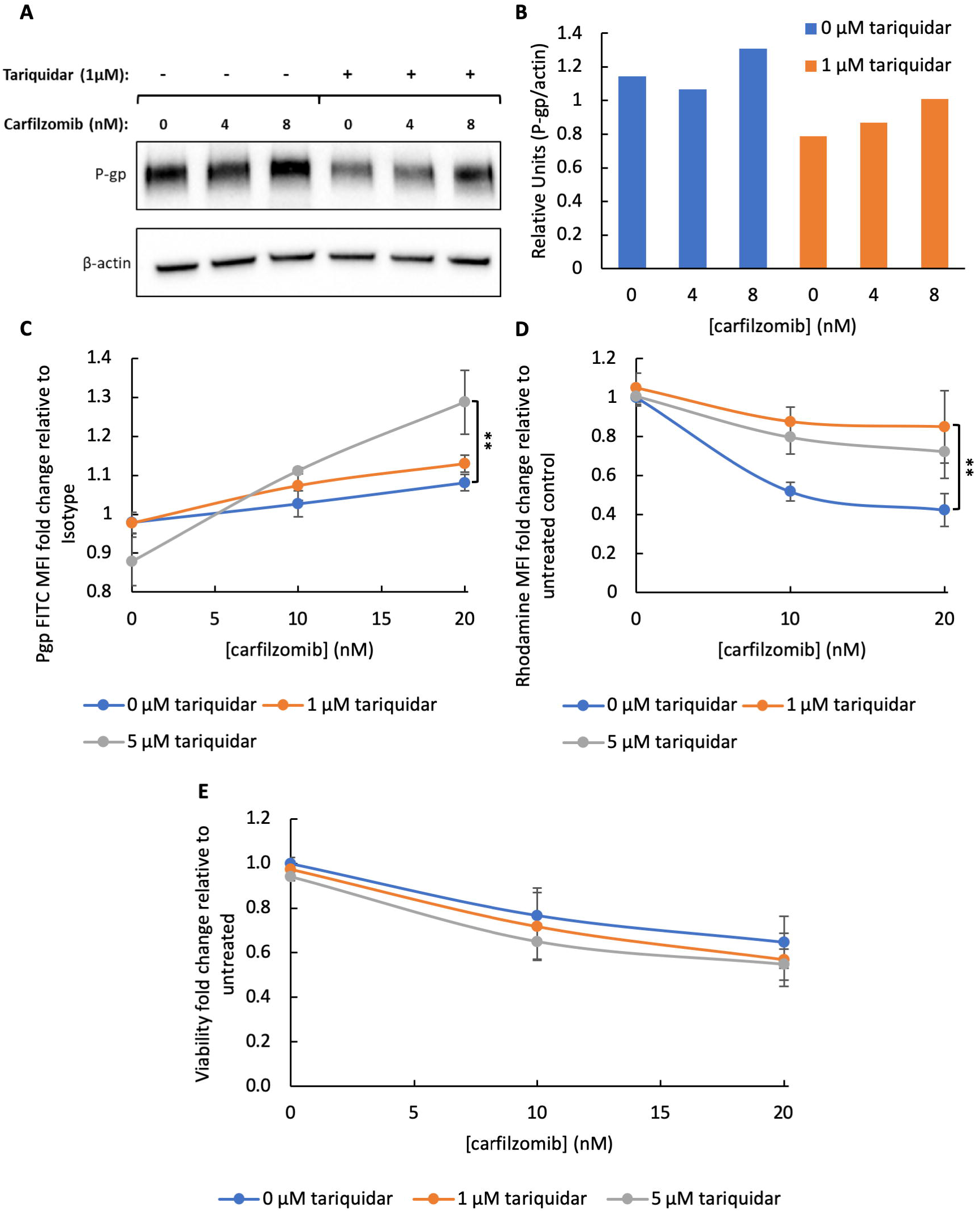
Tariquidar functionally inhibits P-gp but does not synergise with carfilzomib. A) Western blot for KMS-18 cells treated with carfilzomib and 0 or 1 μM tariquidar for 16 hours. B) Densitometric analysis of Western blot displayed in (A), P-gp band intensity relative to respective β-actin loading control. KMS-18 cells were treated with carfilzomib and 0, 1 or 5 μM tariquidar for 24 hours. Surface P-gp expression relative to isotype (C), Rho123 MFI normalised to unstained and relative to untreated (first data point of blue line) controls (D) and viability relative to untreated (E) were analysed by flow cytometry. Flow cytometry data are mean ± standard deviation from three independent experiments. ** p < 0.01.

To determine whether functional inhibition of P-gp can enhance the efficacy of the proteasome inhibitors in the proteasome inhibitor resistant MM cells, RPMI-8226 WT, V10R and C10R cells were next examined. Although all RPMI-8226 cell lines do not exhibit basal P-gp expression, its expression could still be induced by exposure to proteasome inhibitors and thus P-gp may still be of relevance for enhancing the sensitivity of these cells to proteasome inhibitors. However, when RPMI-8226 WT cells were exposed to bortezomib, no induction of P-gp was observed (Fig S1 A and B). Moreover, no changes in Rho123 MFI were seen upon exposure to bortezomib with or without tariquidar (Fig S1 C) and tariquidar did not enhance RPMI-8226 WT cell death in response to bortezomib (Fig S1 D). These data suggest P-gp is not involved in mediating the cytotoxic effects of bortezomib in RPMI-8226 WT cells. The same pattern of results were found when bortezomib resistant RPMI-8226 V10R were cultured with bortezomib (Fig S2) and when RPMI-8226 WT cells (Fig S3) and carfilzomib resistant RPMI-8226 C10R cells (Fig S4) when exposed to carfilzomib. Thus, P-gp does not appear to be involved in mediating bortezomib and carfilzomib resistance in RPMI-8226 cells.

### Inhibiting glucosylceramide synthase

The potent and specific GCS inhibitor eliglustat alone did not alter total P-gp expression in KMS-18 cells, however, 30 μM eliglustat, the highest concentration examined, augmented the increase in P-gp induced by bortezomib (Fig 7A and B). Intriguingly, this combination did not result in increased surface P-gp expression (Fig 7C). P-gp activity as indicated by Rho123 cell efflux did not appear to be affected by eliglustat treatment as the MFI of the dye decreased to the same extent when comparing bortezomib alone to bortezomib plus eliglustat treatment (Fig 7D). These data agree with previous findings demonstrating GCS downregulation by antisense transfection did not influence Rho123 efflux in an adriamycin-resistant breast adenocarcinoma cell line with high P-gp expression (33). The addition of eliglustat did, however, further sensitise KMS-18 cells to bortezomib, as a synergistic reduction in cell viability was seen with bortezomib and 30 μM eliglustat (Fig 7E) (fractional product −0.314 and −0.216 for 4 nM and 10 nM bortezomib, respectively), despite eliglustat being shown to inhibit GCS at concentrations in the low nanomolar range (19, 20).

**Fig 7.**
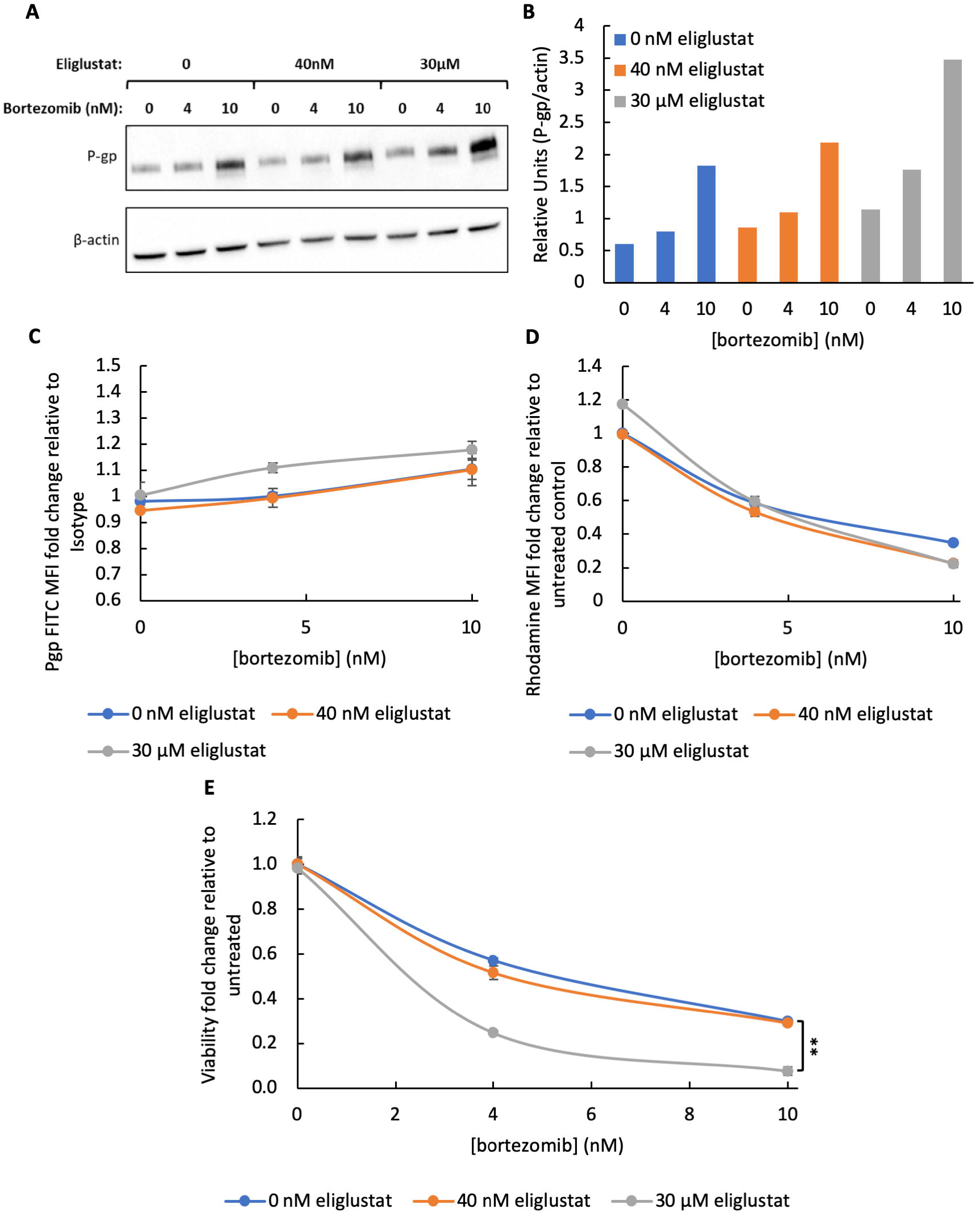
Eliglustat does not inhibit P-gp but synergises with bortezomib. A) Western blot of KMS-18 cells treated with bortezomib and 0, 40 nM or 30 μM eliglustat for 16 hours. B) Densitometric analysis of the Western blot displayed in (A), P-gp band intensity relative to respective β-actin loading control. KMS-18 cells were treated with bortezomib and 0, 40 nM or 30 μM eliglustat for 24 hours. Surface P-gp expression relative to isotype (C), Rho123 MFI normalised to unstained and relative to untreated (first data point of blue line) controls (D) and viability relative to untreated control (E) were analysed by flow cytometry. Flow cytometry data are mean ± standard deviation of duplicate measurements from three independent experiments. ** p < 0.01.

The addition of 30 μM eliglustat to carfilzomib treatment also produced an increase in total P-gp (Fig 8A and B), and to a lesser extent surface P-gp (Fig 8C). Rho123 efflux and thus P-gp activity did not appear to be affected by eliglustat treatment as the MFI of the dye decreased to the same extent when comparing carfilzomib alone to carfilzomib plus eliglustat treatment (Fig 8D). This correlates with the bortezomib and eliglustat data (Fig 7), suggesting inhibition of GCS by eliglustat does not inhibit P-gp activity in KMS-18 cells. The addition of eliglustat did, however, appear to further sensitise the cells to carfilzomib treatment, as a synergistic reduction in cell viability was seen with carfilzomib and 30 μM eliglustat (Fig 8E) (fractional product −0.446 and −0.390 for 10 nM and 20 nM carfilzomib, respectively).

**Fig 8.**
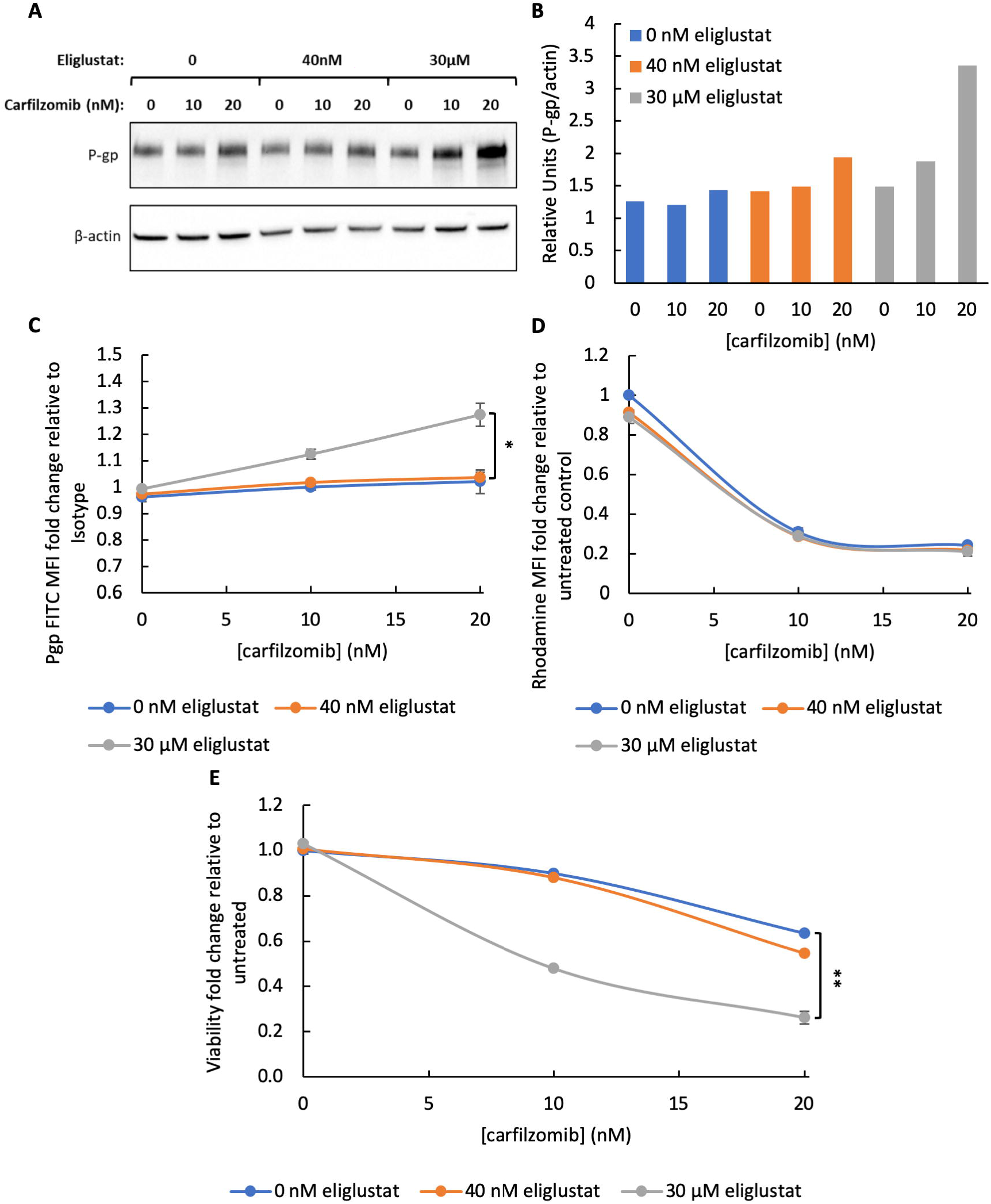
Eliglustat does not inhibit P-gp but synergises with carfilzomib. A) Western blot for KMS-18 cells treated with carfilzomib and 0, 40 nM or 30 μM eliglustat for 16 hours. B) Densitometric analysis of the Western blot displayed in (A), P-gp band intensity relative to respective β-actin loading control. KMS-18 cells were treated with carfilzomib and 0, 40 nM or 30 μM eliglustat for 24 hours. Surface P-gp expression relative to isotype (C), Rho123 MFI normalised to unstained and relative to untreated (first data point of blue line) controls (D) and viability relative to untreated control (E) were analysed by flow cytometry. Flow cytometry data are mean ± standard deviation of duplicate measurements from three independent experiments. *, p < 0.05, ** p < 0.01.

As a comparison to KMS-18 cells, LP-1 cells do not express P-gp and unlike KMS-18 cells, bortezomib treatment does not upregulate P-gp protein expression by Western blot (Fig 9A). Rho123 MFI remains constant with bortezomib alone or in combination with eliglustat, suggesting the dye is being retained in the cell, likely due to the absence of P-gp (Fig 9B). However, LP-1 cells appear to be more sensitive to eliglustat treatment alone at 30 μM compared to KMS-18, as a 20% reduction in viability in LP-1 cells was observed (Fig 9C), compared to no significant change in viability in KMS-18 cells (Figs 7E and 8E). Moreover, a synergistic reduction in cell viability with the combination of drugs was not evident as was seen in KMS-18 cells which do express P-gp (Fig 9C).

**Fig 9.**
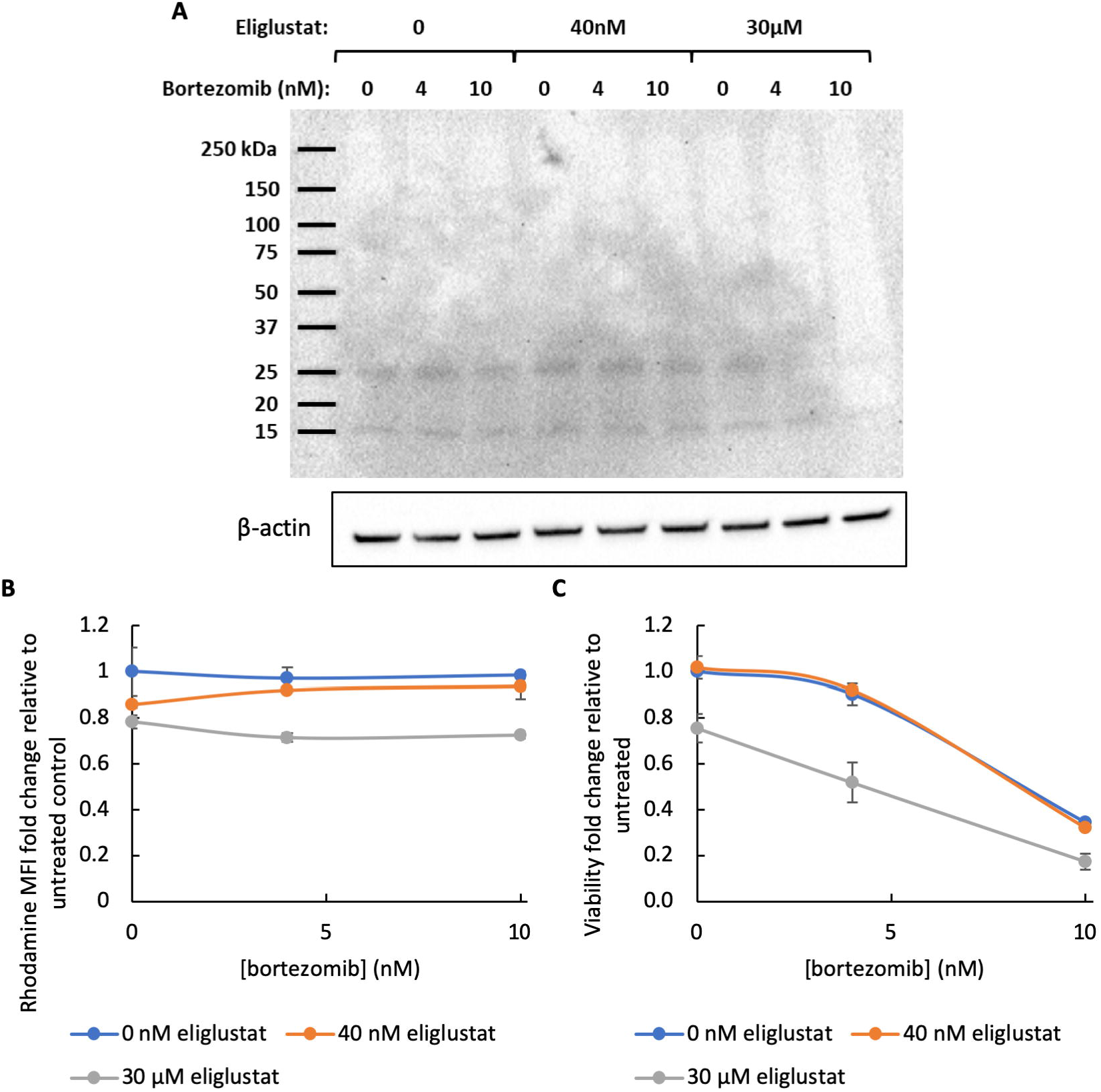
LP-1 cells do not express P-glycoprotein. A) Western blot for LP-1 cells treated with bortezomib and 0, 40 nM or 30 μM eliglustat for 16 hours, β-actin used as a loading control. LP-1 cells were treated with bortezomib and 0, 40 nM or 30 μM eliglustat for 24 hours and the Rho123 MFI normalised to unstained and relative to untreated (first data point of blue line) controls (B) and viability relative to untreated control (C) were analysed by flow cytometry. Flow cytometry data are mean ± standard deviation of duplicate measurements from three independent experiments.

## Discussion

Despite the most well-known and researched function of P-gp being that of drug efflux when located on the plasma membrane, it has also been shown to play a role in drug resistance when situated in intracellular organelles such as lysosomes, mitochondria and nuclei (34–36). This may explain the strong presence of P-gp in KMS-18 whole cell lysates by Western blot whilst flow cytometric analysis demonstrated that it was not expressed to a great extent on the cell surface. Despite these cells not expressing high levels of P-gp on the plasma membrane before or after proteasome inhibitor treatment, they did show a decrease in Rho123 fluorescence after being exposed to these agents, which is indicative of active P-gp. Whilst this may be because the low level of P-gp already on the surface is sufficient to pump out the dye, it could also be related to the greater whole cell P-gp expression observed after proteasome inhibitor treatment and catabolism of the dye in intracellular compartments such as lysosomes on which P-gp is present (37). It has also been suggested that P-gp is regulated by post-translational modifications such phosphorylation and assessing these modifications may be useful to determine whether they have an effect on the activity of P-gp without changing the level of protein on the surface (38).

The other eight human MM cell lines tested, including bortezomib and carfilzomib resistant RPMI-8226 cells, did not express P-gp on the cell surface or in whole cell lysates. Furthermore, its expression was not induced by proteasome inhibition in RPMI-8226 WT, RPMI-8226 V10R, RPMI-8226 C10R, LP-1 and H929 cells, indicated by a lack of protein band at 170 kDa (H929 Western blot displayed in Fig S5). The Rho123 efflux studies confirm this as the dye was retained in RPMI-8226 and LP-1 cells treated with bortezomib and/or carfilzomib, a response similar to that observed for KMS-18 cells when P-gp was inhibited (Figs 5 and 6). Our RNA-seq data also support this, with minimal P-gp gene expression found in LP-1 cells treated with proteasome inhibitors, therefore this cell line and the remaining MM cell lines, except for the RPMI-8226 series, were not used in further P-gp experiments. These data support the published findings that most MM cell lines have low P-gp expression, making it difficult to test its contribution to proteasome inhibitor cellular efflux (12). Unfortunately, the doxorubicin resistant MM cell line (RPMI-8226/Dox40) that has been shown to have high P-gp expression could not be obtained and would be a useful positive control for experiments assessing P-gp expression and function (11, 39).

Tariquidar is a functional inhibitor of P-gp, blocking its ATPase activity, and as expected it did not directly affect P-gp expression (8). Tariquidar was shown to effectively inhibit P-gp in MM cells at a concentration of 5 μM (8) and 1 μM in other cancers (32), so these concentrations were chosen for further investigation. Both 1 μM and 5 μM tariquidar resulted in a decrease in P-gp activity when combined with bortezomib, as shown by an accumulation of Rho123 in the KMS-18 cells. Conversely, bortezomib treatment alone resulted in an increase in P-gp activity in these cells. These data contradict a previous report where after 24 hours in culture with bortezomib, RPMI-8226/Dox40 MM cells had a reduction in P-gp activity shown by an accumulation of Rho123 in the cells and a reduction of P-gp protein expression by Western blot (11). Despite tariquidar inhibiting P-gp, it did not further sensitise the cells to bortezomib. This is consistent with previous studies showing bortezomib is not a substrate of P-gp (6, 13–15), however, contradicts research that showed inhibition of P-gp does improve the efficacy of bortezomib in MM cell lines (11, 12).

Carfilzomib requires higher concentrations compared to bortezomib to produce an increase in total cell P-gp expression by Western blot in KMS-18 cells. Similar to bortezomib, it does not induce P-gp expression on the cell surface by flow cytometry, however, P-gp activity does increase with increasing concentrations of carfilzomib as seen by decreased Rho123 fluorescence that can be reversed by tariquidar. The combination of 5 μM tariquidar with 20 nM carfilzomib did cause a very small increase in surface P-gp which may be to compensate for P-gp inhibition. Despite the literature suggesting that carfilzomib is a substrate of P-gp (6–9), inhibition of P-gp by tariquidar did not further sensitise KMS-18 cells to carfilzomib treatment.

Importantly, RPMI-8226 cells resistant to bortezomib or carfilzomib, whilst lacking basal P-gp expression, did not increase its expression in response to the proteasome inhibitors nor did P-gp inhibition with tariquidar in these cells enhance proteasome inhibitor cytotoxicity. Taken together, these data demonstrate that P-gp inhibition does not improve the efficacy of bortezomib or carfilzomib, suggesting drug efflux by P-gp may not be a mechanism underpinning proteasome inhibitor resistance in MM.

Despite another research group (8) showing 5 μM tariquidar was sufficient to further sensitise MM.1S MM cells to 5 nM bortezomib and carfilzomib, this concentration of tariquidar did not enhance proteasome inhibitor induced cell death in KMS-18 cells. Using a higher tariquidar concentration and longer incubation period may be valuable to confirm these results in KMS-18 MM cells. It is important to note that whilst no changes in cell viability were observed, changes in cell cycle and proliferation were not examined. It is entirely possible that the combination of tariquidar with a proteasome inhibitor mediates changes in cell proliferation rather than cell viability and this could be assessed in the future.

A link between glucosylceramides and P-gp has been established but is not well understood. Genetic or chemical inhibition of GCS which converts ceramide to glucosylceramide has been shown to downregulate P-gp expression (17) and so these findings led us to use the potent and specific GCS inhibitor eliglustat as a way to indirectly inhibit P-gp. Interestingly, eliglustat did not functionally inhibit P-gp as there were no significant changes in Rho123 fluorescence before or after combining bortezomib/carfilzomib with eliglustat, thereby not supporting the hypothesis that GCS inhibition downregulates P-gp expression and/or activity, at least in MM cells. Eliglustat did, however, produce synergistic cell death when combined with bortezomib and carfilzomib but only at a high concentration (30 μM) despite only needing a concentration in the low nanomolar range to inhibit GCS enzymatic activity (19, 20). This high concentration of eliglustat and the fact that it did not inhibit the activity of P-gp suggests off target effects may be causing this synergistic reduction in cell viability. One research group showed that downregulation of GCS by siRNA-mediated knockdown slowed the clearance of ceramide which is pro-apoptotic, and so an accumulation of ceramide may be contributing to the increase in cell death observed in cells treated with 30 μM eliglustat and a proteasome inhibitor (17). Such a high concentration of eliglustat is likely not clinically achievable so these data suggest eliglustat may not be an effective drug to further sensitise MM cells to proteasome inhibition.

P-gp plays a role in chemotherapy resistance in a number of cancers, however its role in contributing to proteasome inhibitor efficacy in MM is less well studied with conflicting reports in the literature. Our data presented herein suggests that P-gp is poorly expressed in MM cells, including those resistant to proteasome inhibitors, and that inhibition of P-gp does not enhance proteasome inhibitor efficacy. Thus, a strategy to either increase the efficacy of proteasome inhibitors or reverse resistance to these agents in MM via inhibition of P-gp may not be worthy of further exploration.

## Supporting information

Raw Western Blot Images

## Acknowledgements

The authors would like to acknowledge Dr Giles Best from the Flinders University Flow Cytometry Facility for providing support with the flow cytometric assays used for this work.

## Supporting Information

**Fig S5. H929 MM cells do not express P-gp.** Western blot probed for P-gp (170 kDa) in H929 cells treated with bortezomib ± tariquidar for 16 hours, showing H929 cells do not express P-gp.

**Fig S6. Raw Western blot images.**

