## Supplementary material for "Inhibition of p-glycoprotein does not increase the efficacy of proteasome inhibitors in multiple myeloma cells": Raw Western Blot Images

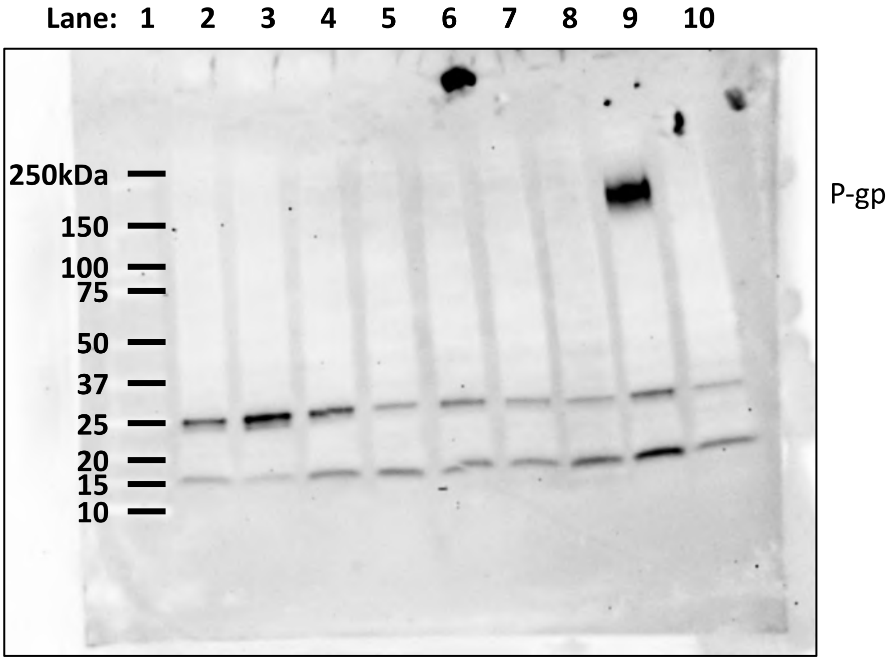


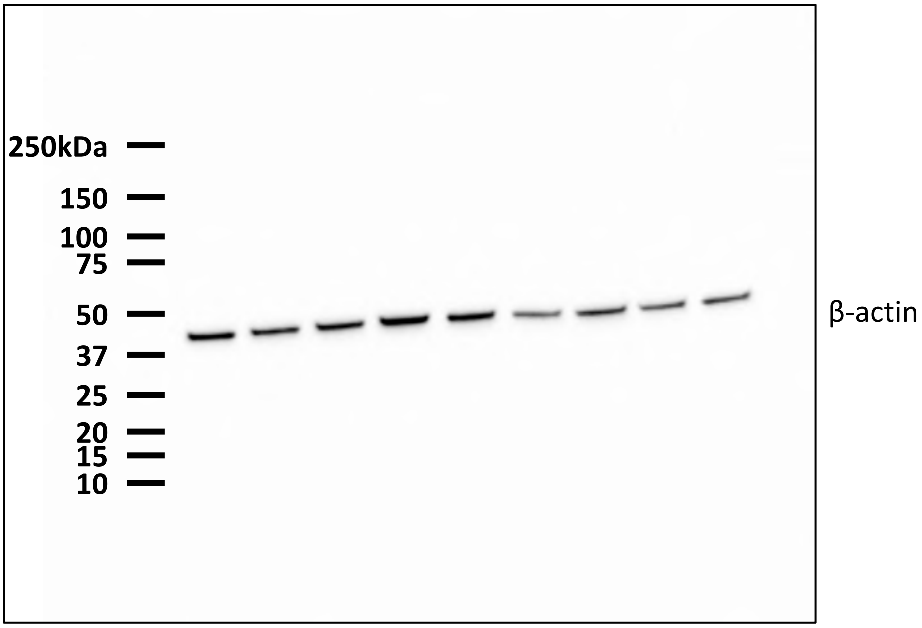


Western blot from Figure 2. Lanes were loaded in numerical order. Lanes 1-10 are; protein standards, RPMI-8226 WT, RPMI-8226 C10R, RPMI-8226 V10R, MM.1S, MM.1R, LP-1, KMS-11, KMS-18 and H929, respectively. Images were obtained on the ChemiDoc MP (Bio-Rad) using the Chemi High Sensitivity blot protocol for an appropriate exposure time.

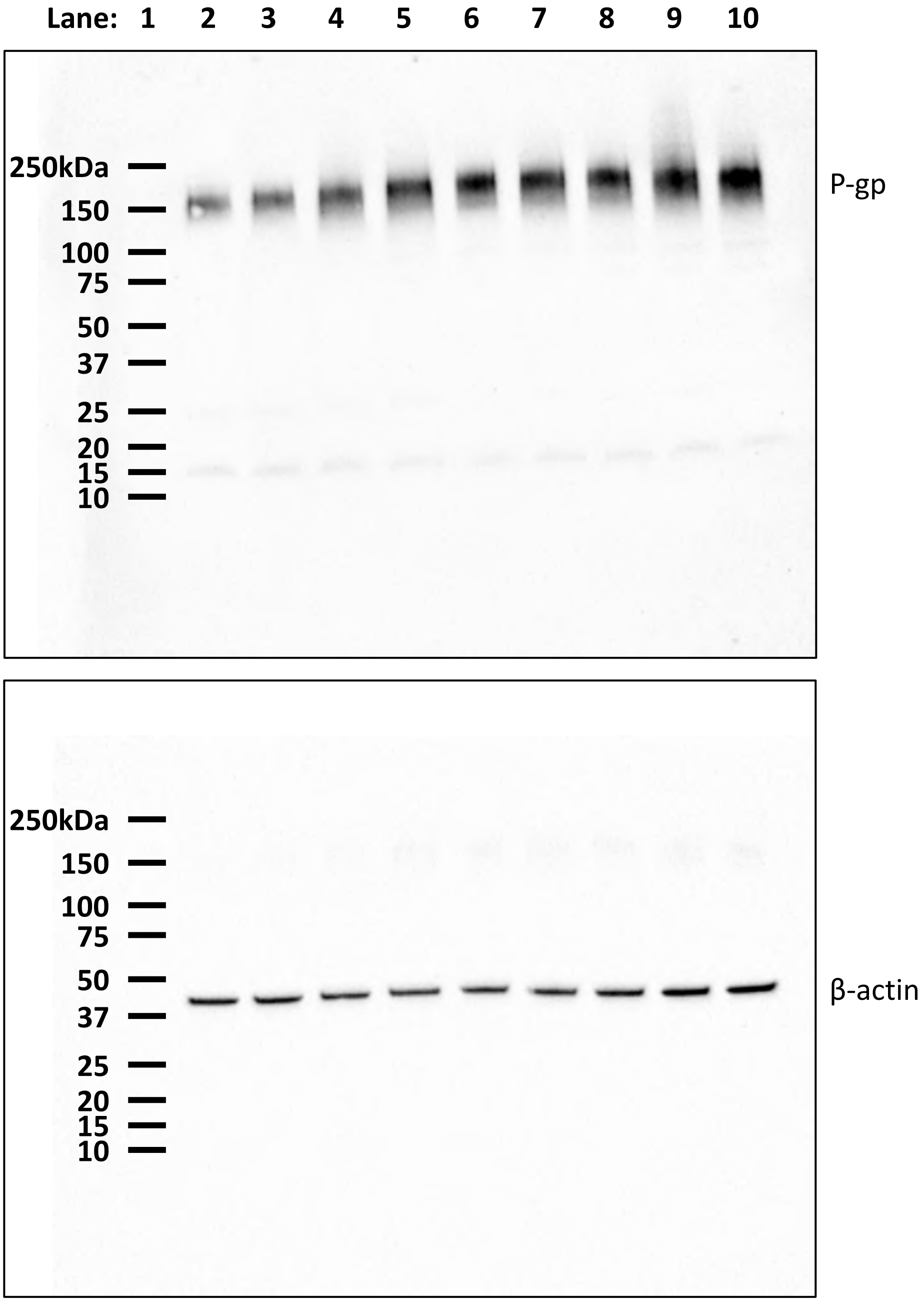


Western blot from Figure 3e. Lanes were loaded in numerical order. Lanes 1-10 are; protein standards, 0, 5, 10, 15, 20, 25, 30, 25 and 40 nM bortezomib respectively. Images were obtained on the ChemiDoc MP (Bio-Rad) using the Chemi High Sensitivity blot protocol for an appropriate exposure time.

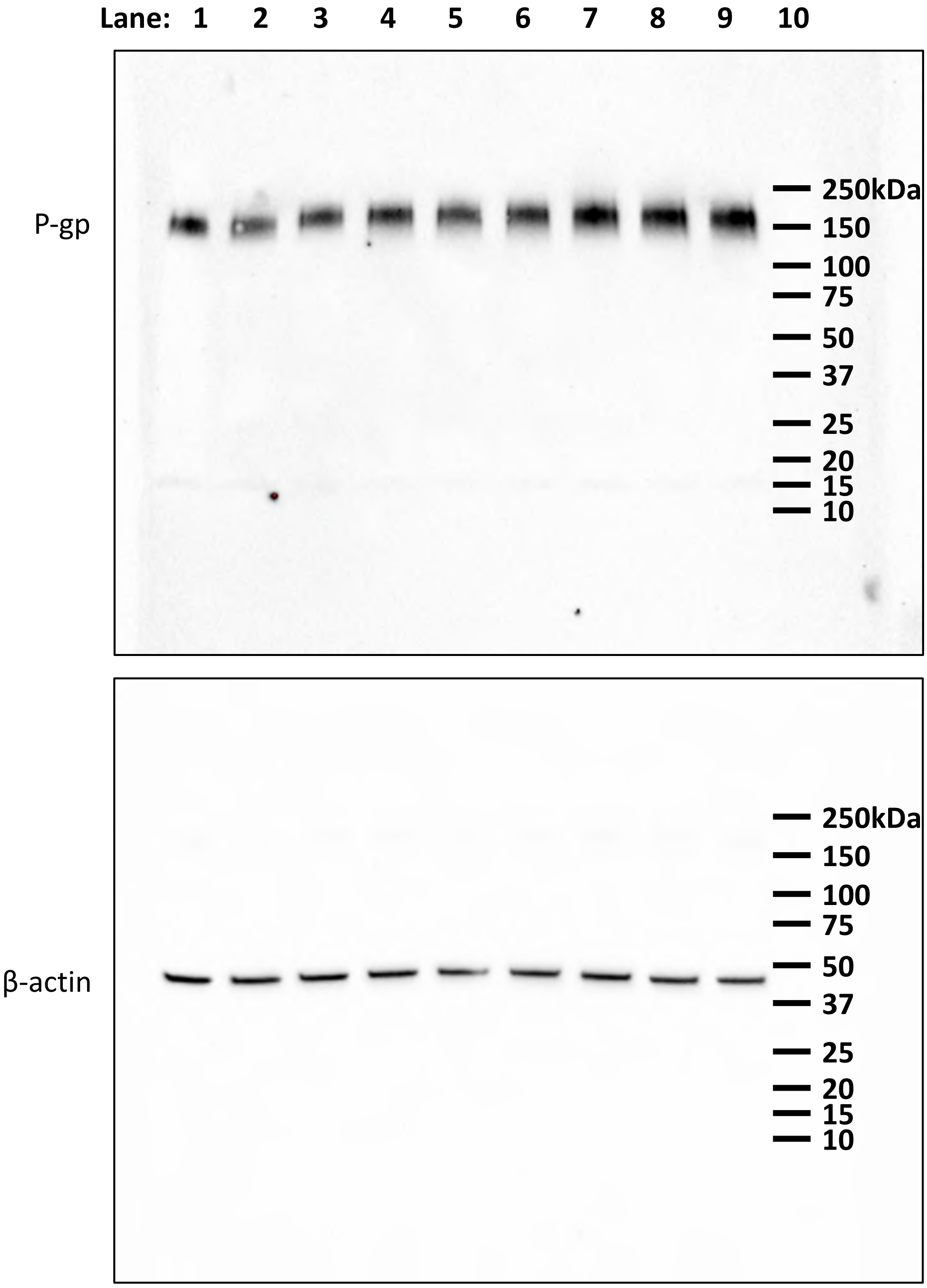


Western blot from Figure 3f. Lane 10 was loaded first and lanes 1-9 were loaded in numerical order. Lanes 1-10 are; 0, 5, 10, 15, 20, 25, 30, 25 and 40 nM carfilzomib and protein standards, respectively. Images were obtained on the ChemiDoc MP (Bio-Rad) using the Chemi High Sensitivity blot protocol for an appropriate exposure time.

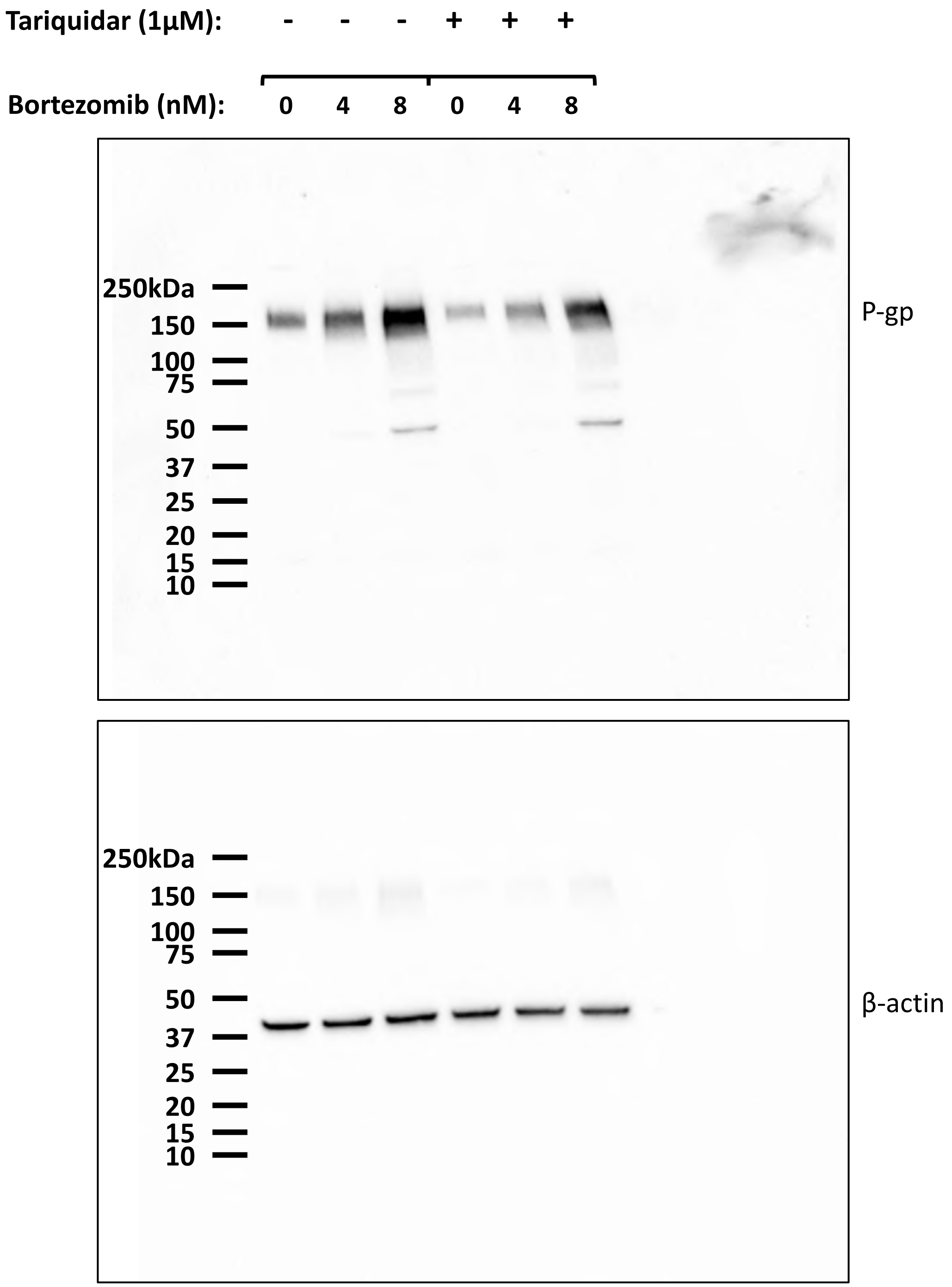


Western blot from Figure 5. Lanes were loaded from left to right. Images were obtained on the ChemiDoc MP (Bio-Rad) using the Chemi High Sensitivity blot protocol for an appropriate exposure time.

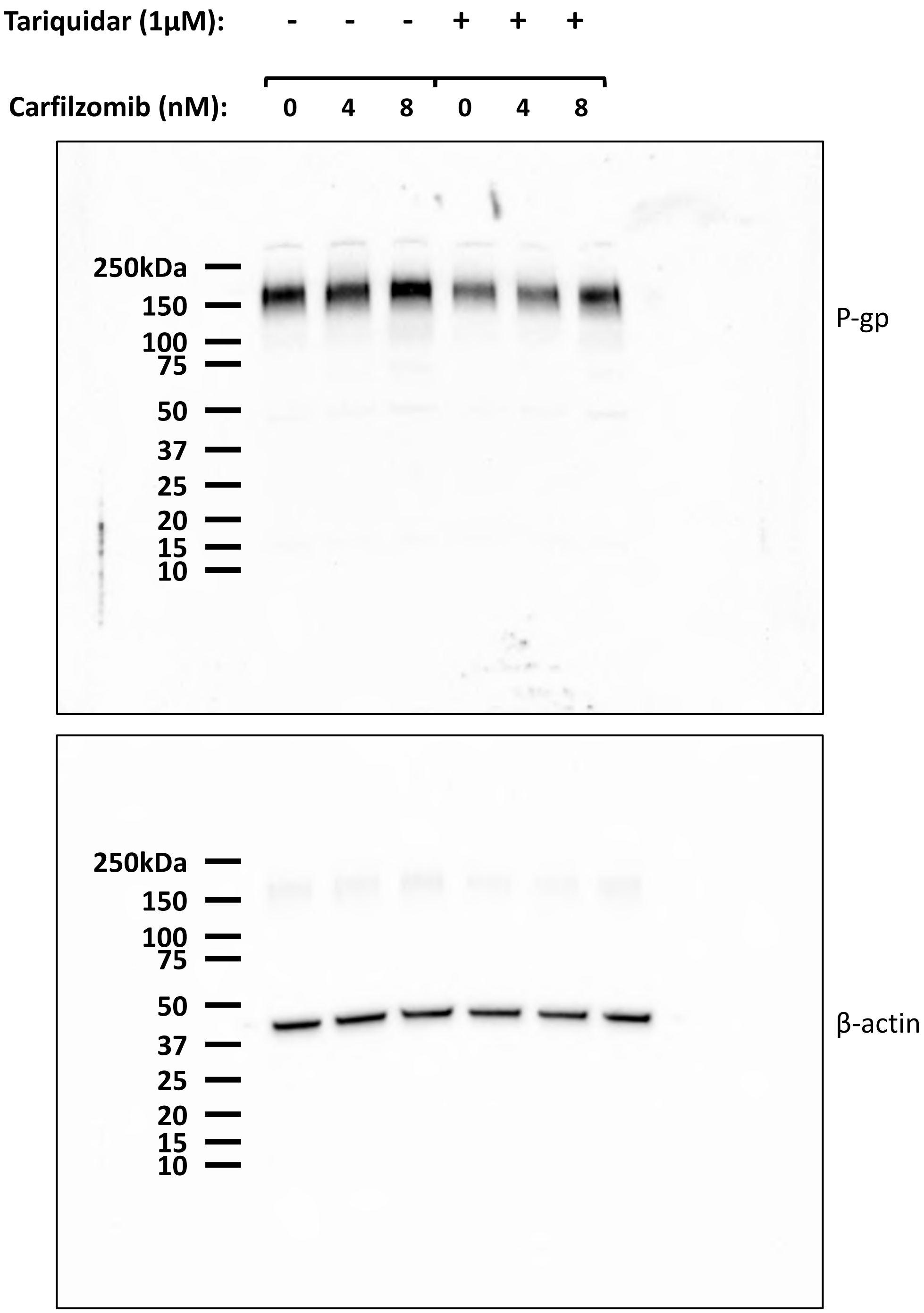


Western blot from Figure 6. Lanes were loaded from left to right. Images were obtained on the ChemiDoc MP (Bio-Rad) using the Chemi High Sensitivity blot protocol for an appropriate exposure time

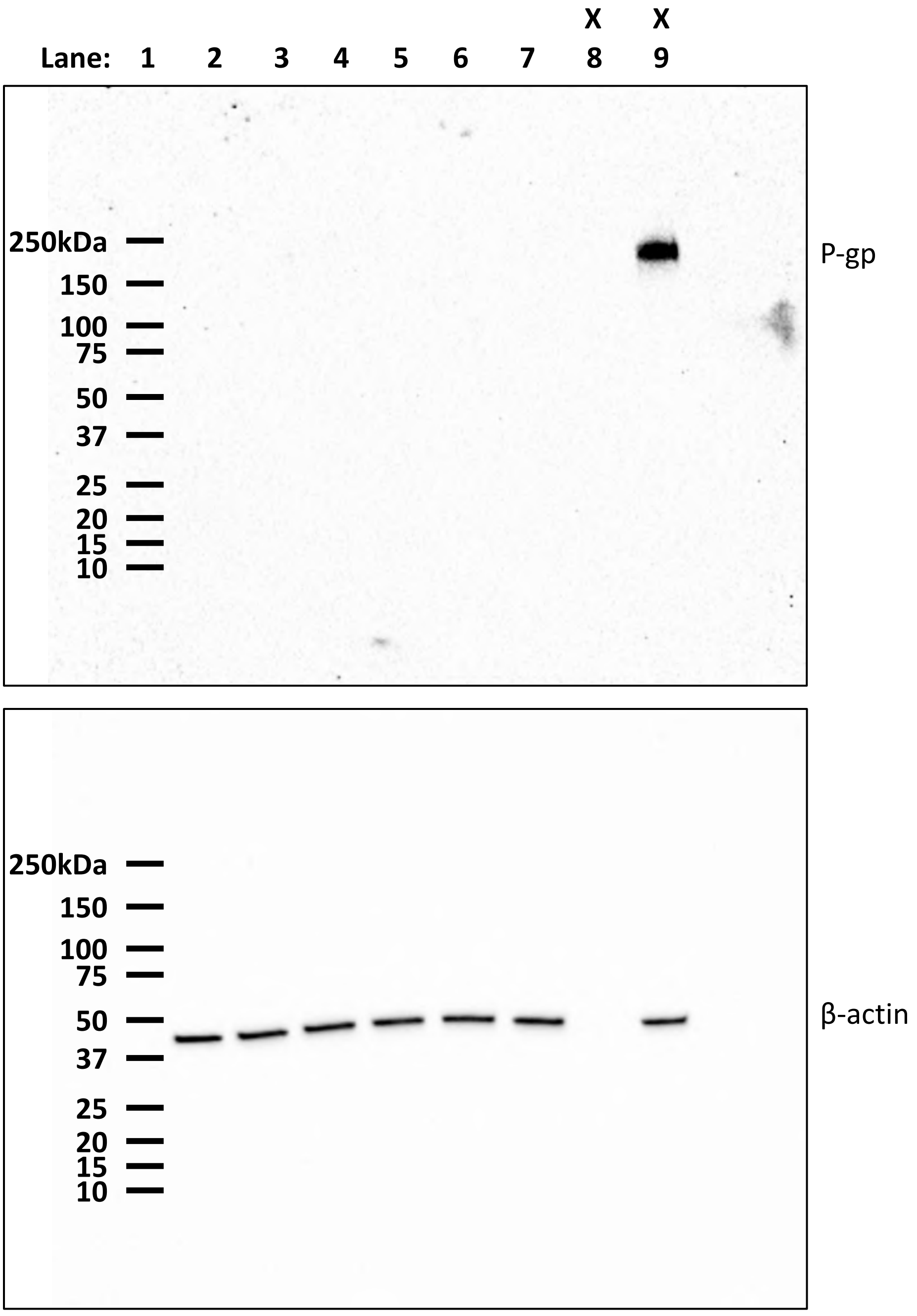


Western blot from Figure 7. Lanes were loaded in numerical order. Lane 1 is protein standards, lanes 2-4 are 0, 4 and 10 nM bortezomib, lanes 5-7 are 0, 4 and 10 nM bortezomib plus 1 μM tariquidar, lane 8 is blank and lane 9 is an untreated KMS-18 positive control. Images were obtained on the ChemiDoc MP (Bio-Rad) using the Chemi High Sensitivity blot protocol for an appropriate exposure time.

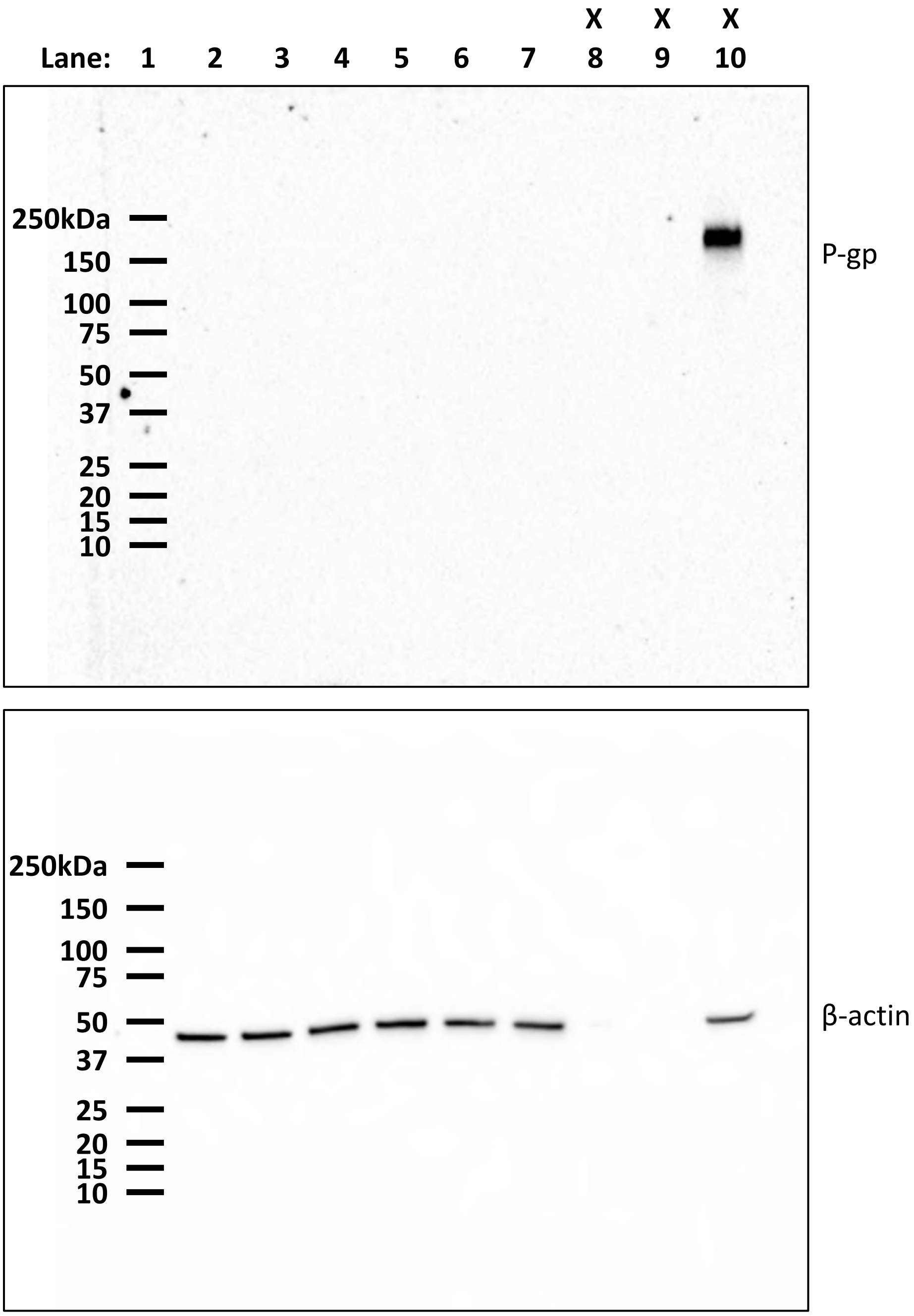


Western blot from Figure 8. Lanes were loaded in numerical order. Lane 1 is protein standards, lanes 2-4 are 0, 20 and 80 nM bortezomib, lanes 5-7 are 0, 20 and 80 nM bortezomib plus 1 μM tariquidar, lanes 8 and 9 are blank and lane 10 is an untreated KMS-18 positive control. Images were obtained on the ChemiDoc MP (Bio-Rad) using the Chemi High Sensitivity blot protocol for an appropriate exposure time.

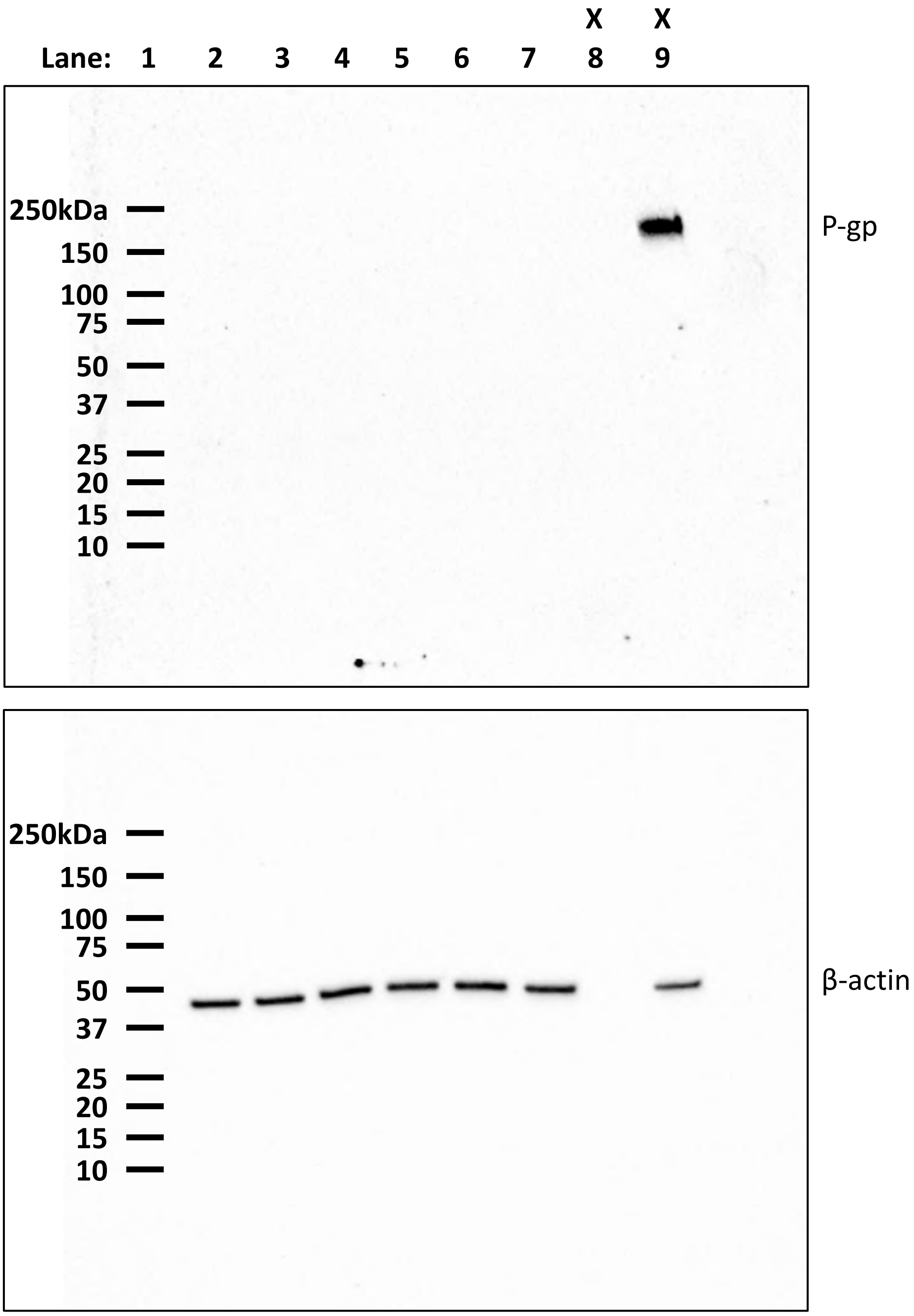


Western blot from Figure 9. Lanes were loaded in numerical order. Lane 1 is protein standards, lanes 2-4 are 0, 2 and 8 nM carfilzomib, lanes 5-7 are 0, 2 and 8 nM carfilzomib plus 1 µM tariquidar, lane 8 is blank and lane 9 is an untreated KMS-18 positive control. Images were obtained on the ChemiDoc MP (Bio-Rad) using the Chemi High Sensitivity blot protocol for an appropriate exposure time.

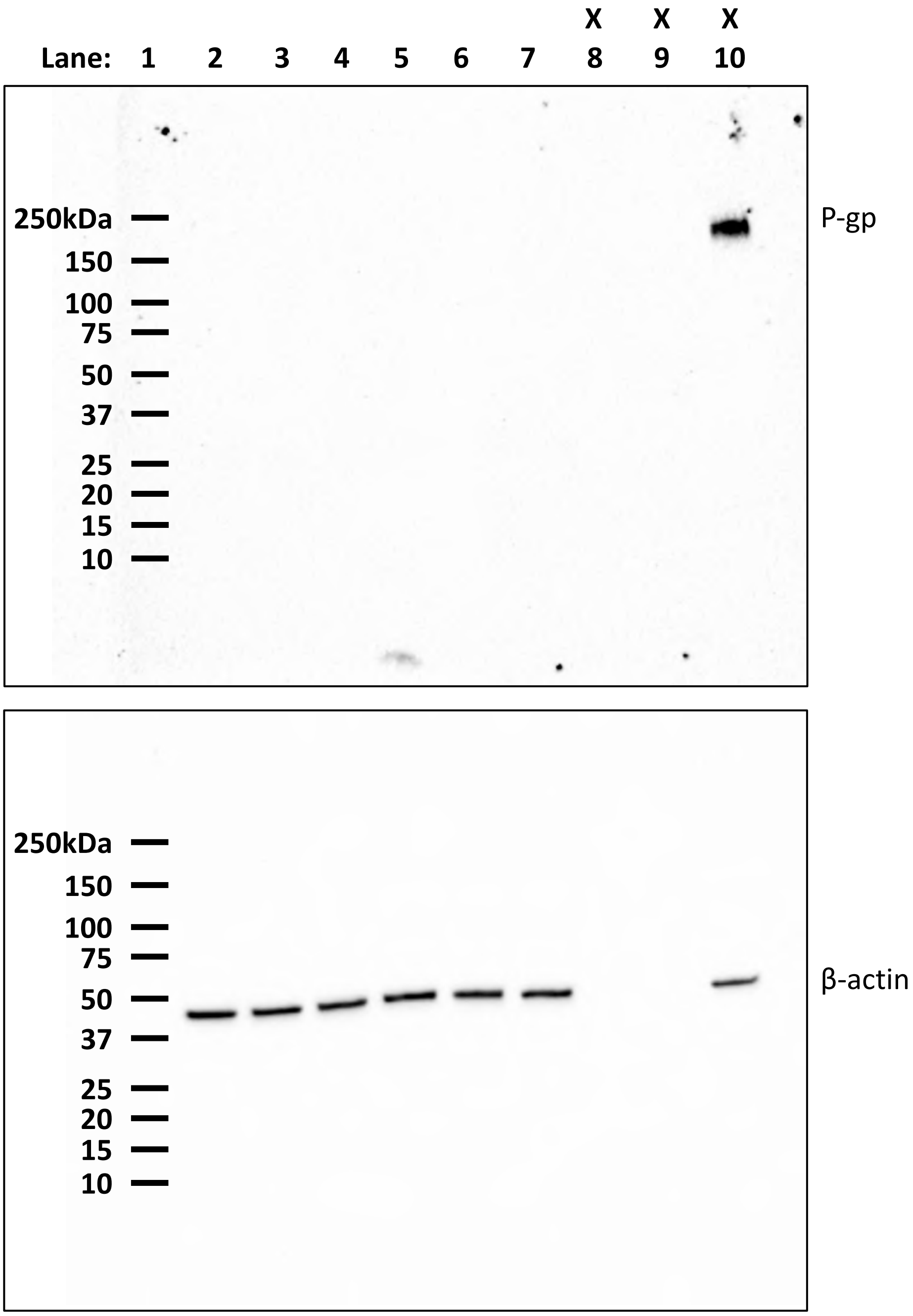


Western blot from Figure 10. Lanes were loaded in numerical order. Lane 1 is protein standards, lanes 2-4 are 0, 10 and 20 nM carfilzomib, lanes 5-7 are 0, 10 and 20 nM carfilzomib plus 1 μM tariquidar, lanes 8 and 9 are blank and lane 10 is an untreated KMS-18 positive control. Images were obtained on the ChemiDoc MP (Bio-Rad) using the Chemi High Sensitivity blot protocol for an appropriate exposure time.

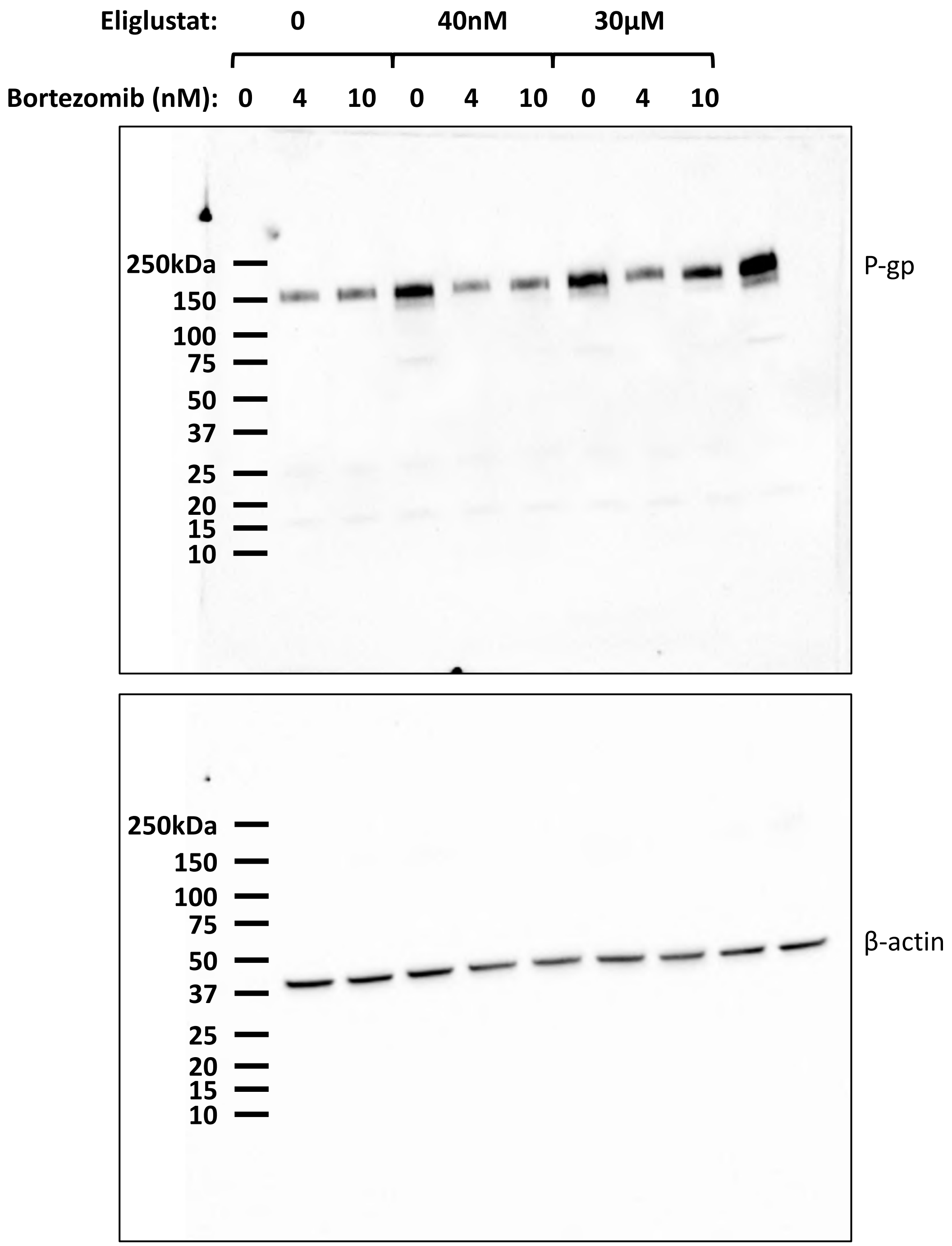


Western blot from Figure 11. Lanes were loaded from left to right. Images were obtained on the ChemiDoc MP (Bio-Rad) using the Chemi High Sensitivity blot protocol for an appropriate exposure time.

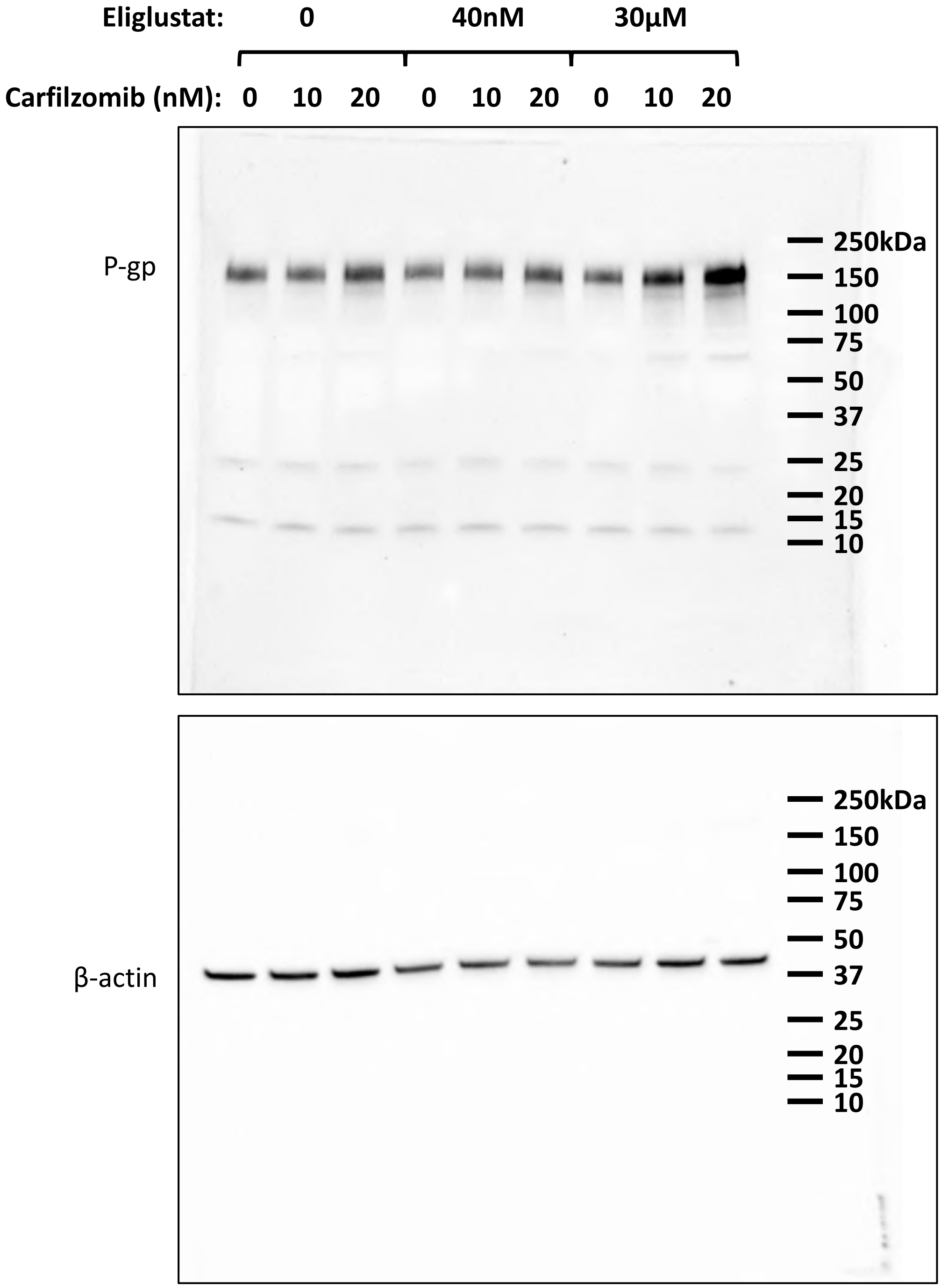


Western blot from Figure 12. Molecular weight ladder was loaded first and then the sample lanes were loaded from left to right. Images were obtained on the ChemiDoc MP (Bio-Rad) using the Chemi High Sensitivity blot protocol for an appropriate exposure time.

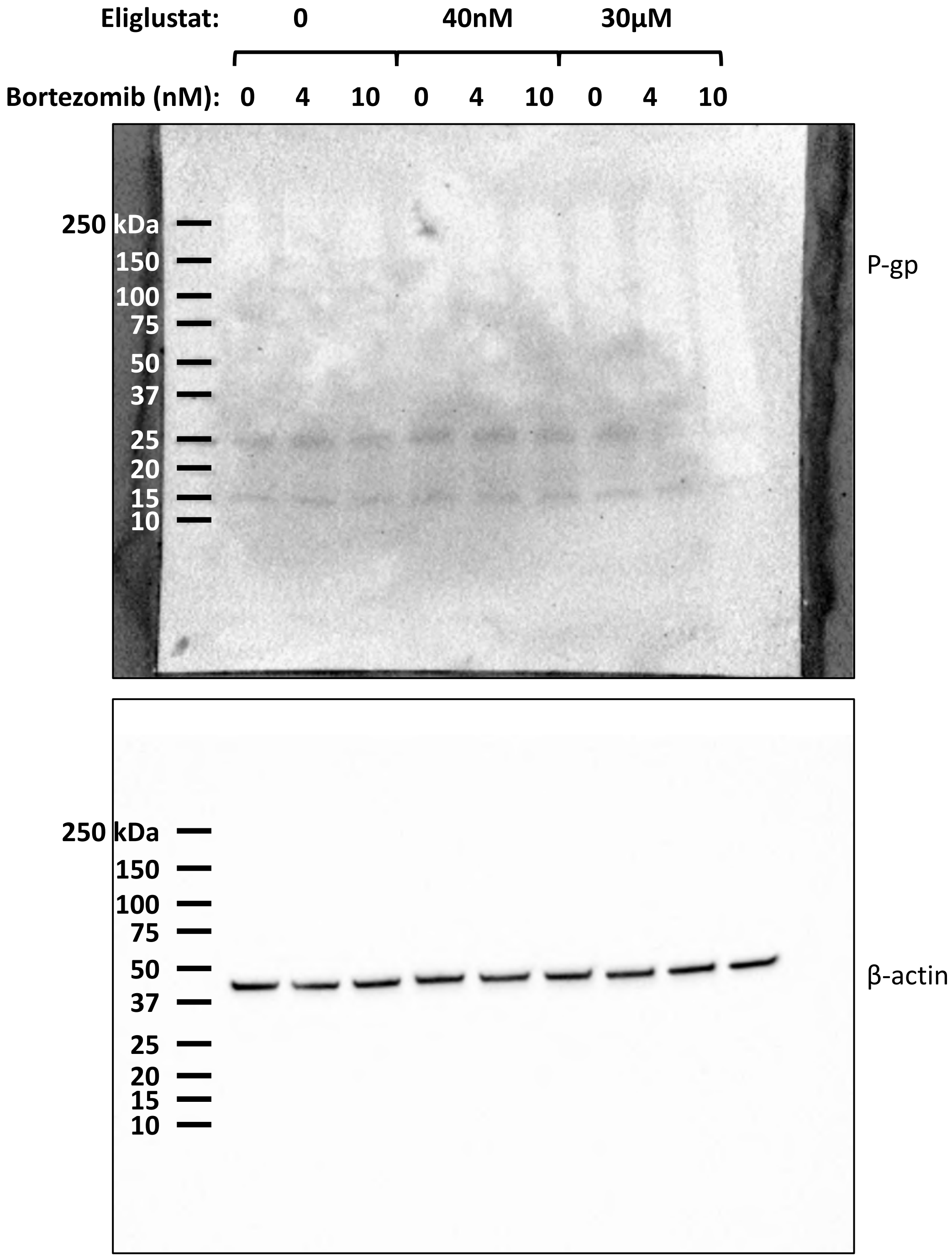


Western blot from Figure 13. Lanes were loaded from left to right. Images were obtained on the ChemiDoc MP (Bio-Rad) using the Chemi High Sensitivity blot protocol for an appropriate exposure time.
