## Supplementary Figure Legends for "Inhibition of p-glycoprotein does not increase the efficacy of proteasome inhibitors in multiple myeloma cells"

**Supplementary Data**

**Fig S1. Tariquidar has no effect on bortezomib efficacy in wild type RPMI-8226 MM cells**. A) Western blot for RPMI-8226 WT cells treated with bortezomib and 0 or 1 µM tariquidar for 16 hours. RPMI-8226 WT cells were treated with bortezomib and 0, 1 or 5 µM tariquidar for 24 hours. Surface P-gp expression relative to isotype (B), Rho123 MFI normalised to unstained and relative to untreated (first data point of blue line) controls (C) and viability relative to untreated control (D) were analysed by flow cytometry. Flow cytometry data are mean ± standard deviation of duplicate measurements from three independent experiments.

**Fig S2. Tariquidar has no effect on bortezomib efficacy in bortezomib resistant RPMI-8226 MM cells**. A) Western blot for RPMI-8226 V10R cells treated with bortezomib and 0 or 1 µM tariquidar for 16 hours. RPMI-8226 V10R cells were treated with bortezomib and 0, 1 or 5 µM tariquidar for 24 hours. Surface P-gp expression relative to isotype (B), Rho123 MFI normalised to unstained and relative to untreated (first data point of blue line) controls (C) and viability relative to untreated control (D) were analysed by flow cytometry. Flow cytometry data are mean ± standard deviation of duplicate measurements from three independent expeirments.

**Fig S3. Tariquidar has no effect on carfilzomib efficacy in wild type RPMI-8226 MM cells**. A) Western blot for RPMI-8226 WT cells treated with carfilzomib and 0 or 1 µM tariquidar for 16 hours. RPMI-8226 WT cells were treated with carfilzomib and 0, 1 or 5 µM tariquidar for 24 hours. Surface P-gp expression relative to isotype (B), Rho123 MFI normalised to unstained and relative to untreated (first data point of blue line) controls (C) and viability relative to untreated control (D) were analysed by flow cytometry. Flow cytometry data are mean ± standard deviation of duplicate measurements from three independent expeirments.

**Fig S4. Tariquidar has no effect on carfilzomib efficacy in carfilzomib resistant RPMI-8226 MM cells**. A) Western blot for RPMI-8226 C10R cells treated with carfilzomib and 0 or 1 µM tariquidar for 16 hours. RPMI-8226 C10R cells were treated with carfilzomib and 0, 1 or 5 µM tariquidar for 24 hours. Surface P-gp expression relative to isotype (B), Rho123 MFI normalised to unstained and relative to untreated (first data point of blue line) controls (C) and viability relative to untreated control (D) were analysed by flow cytometry. Flow cytometry data are mean ± standard deviation of duplicate measurements from three independent experiments.

**Fig S5. P-gp expression in H929 MM cells with bortezomib and tariquidar**. A) Western blot for H929 cells treated with bortezomib and 0 or 1 µM tariquidar for 16 hours.
