## Supplementary figures and images for "Inhibition of p-glycoprotein does not increase the efficacy of proteasome inhibitors in multiple myeloma cells"

### Supplementary Figure 1

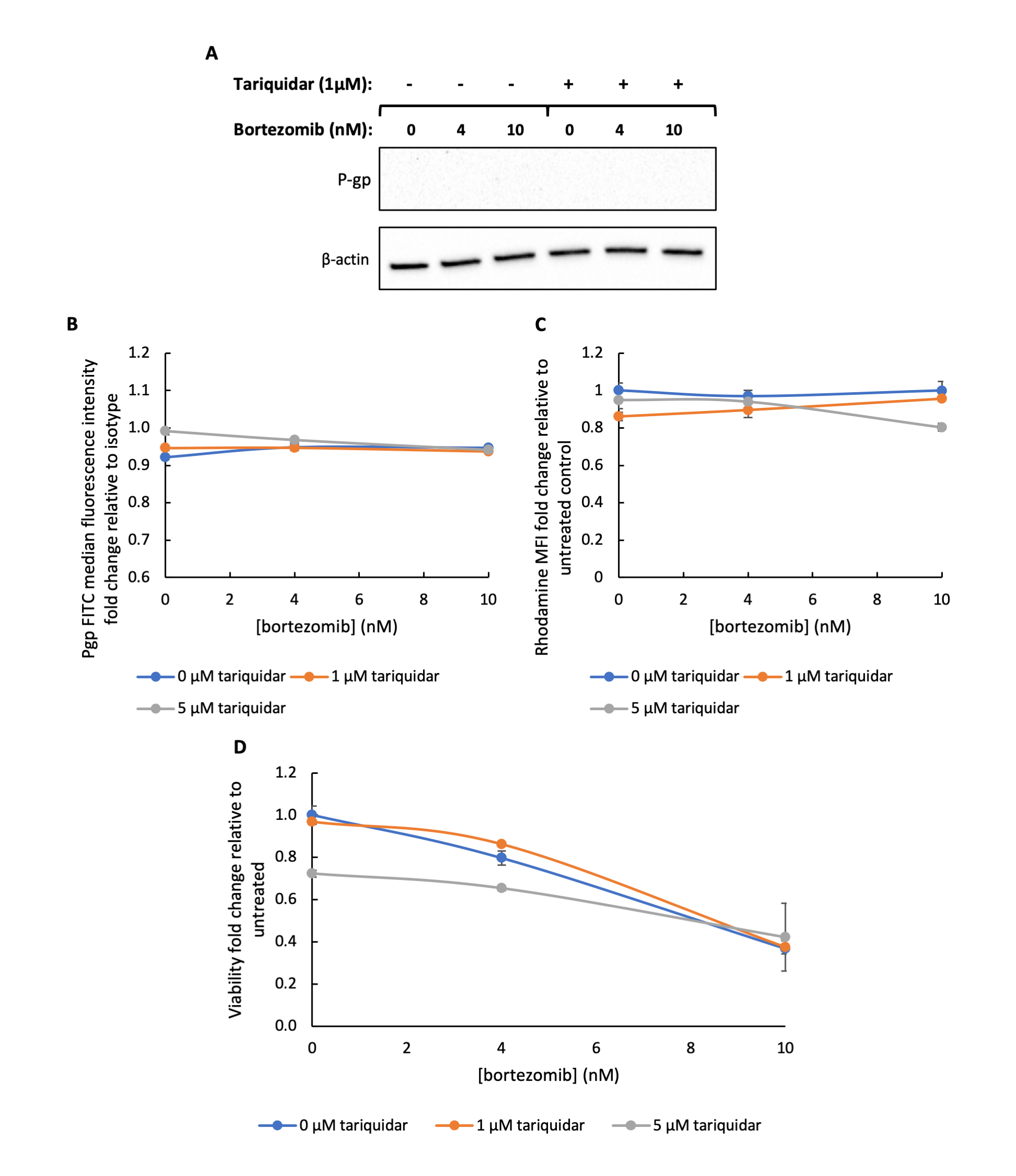

### Supplementary Figure 2

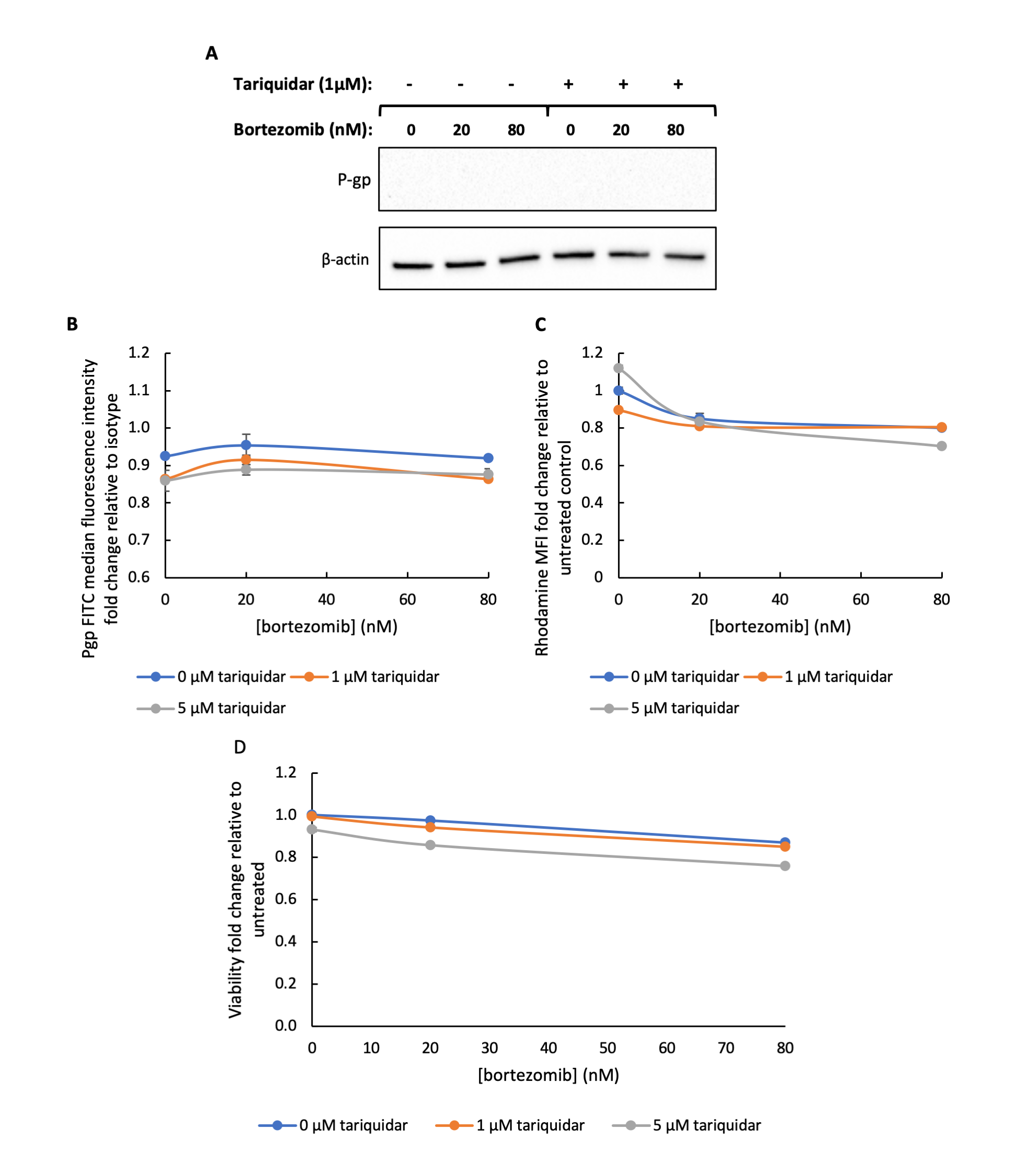

### Supplementary Figure 3

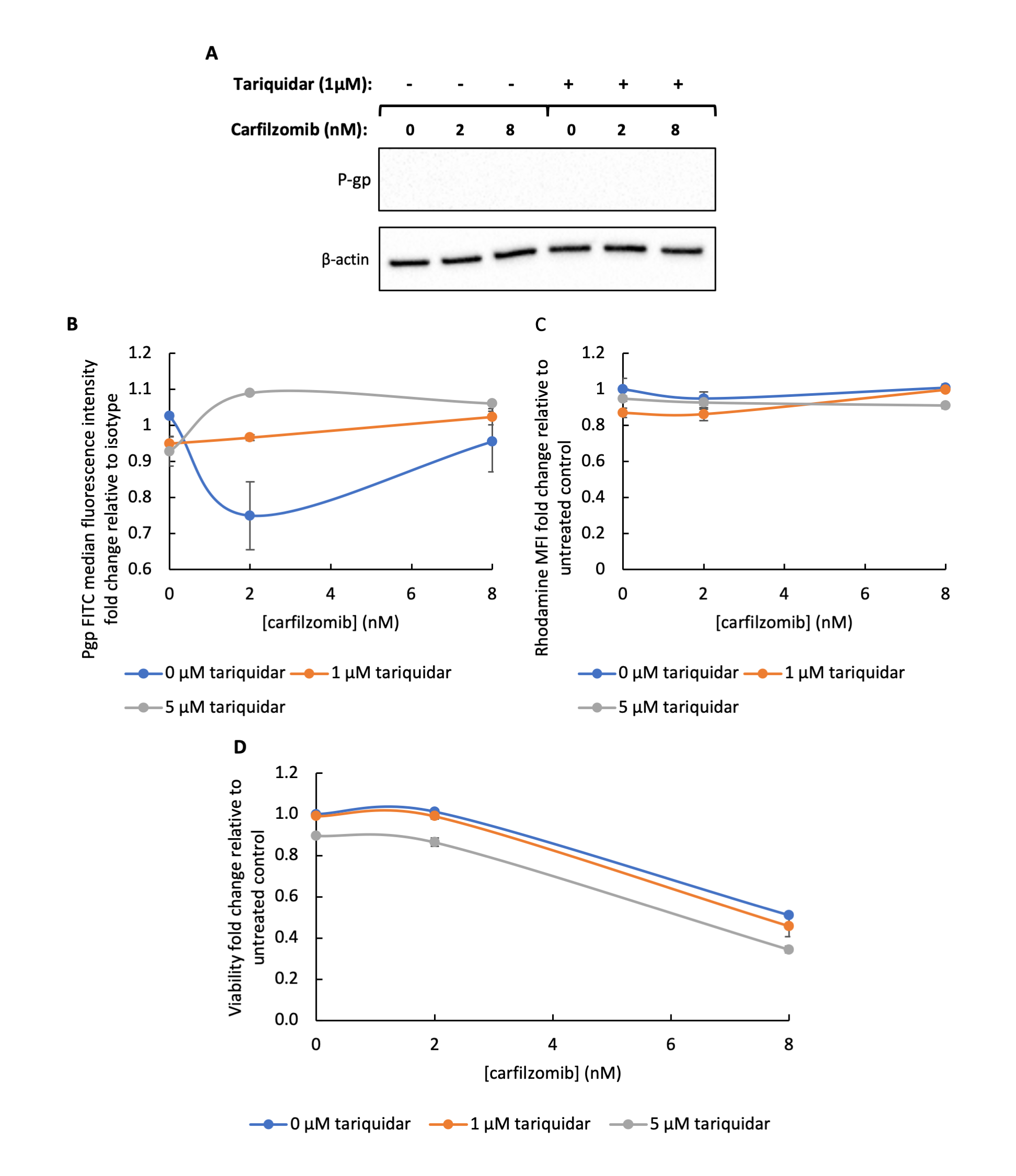

### Supplementary Figure 4

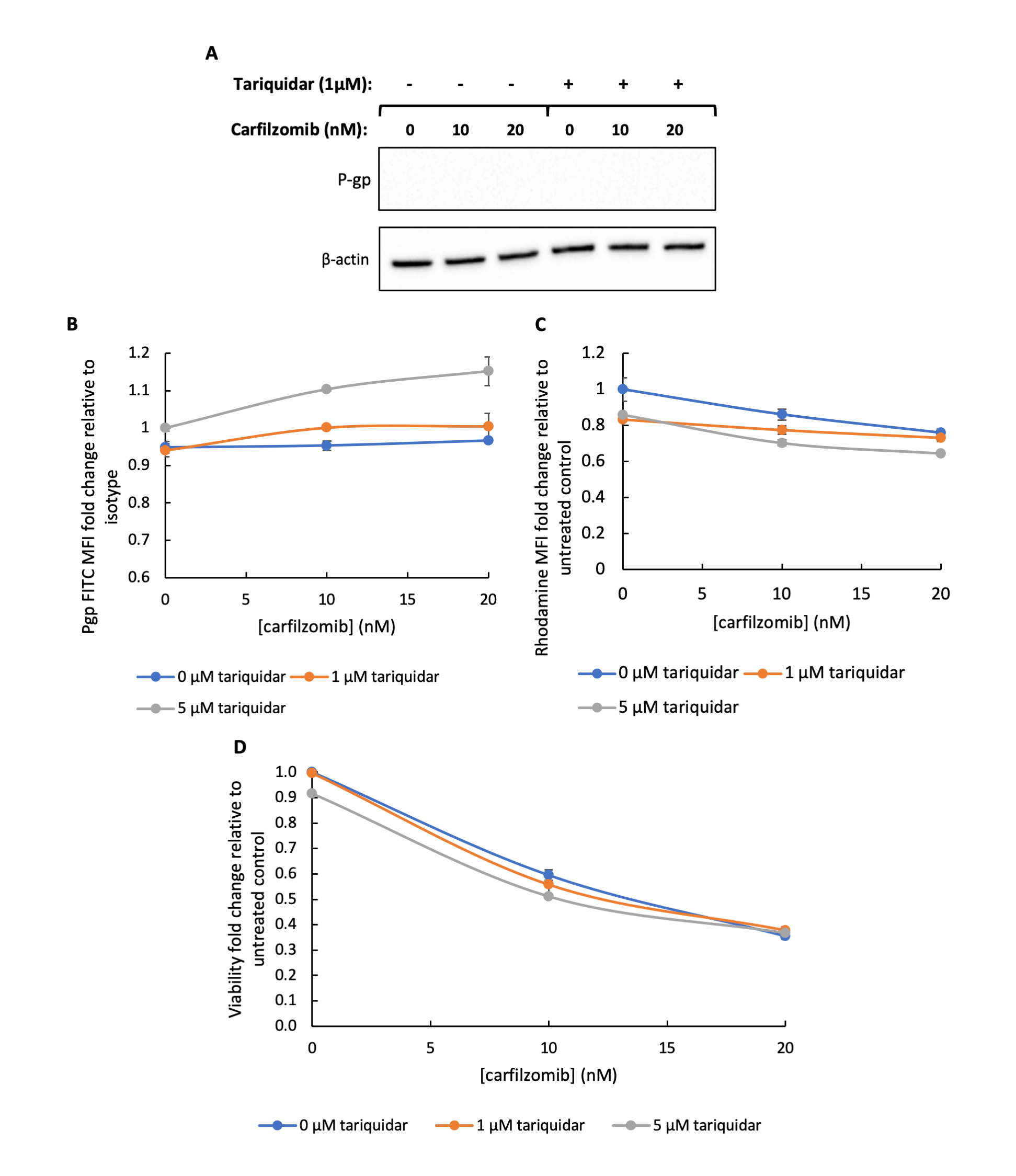

### Supplementary Figure 5

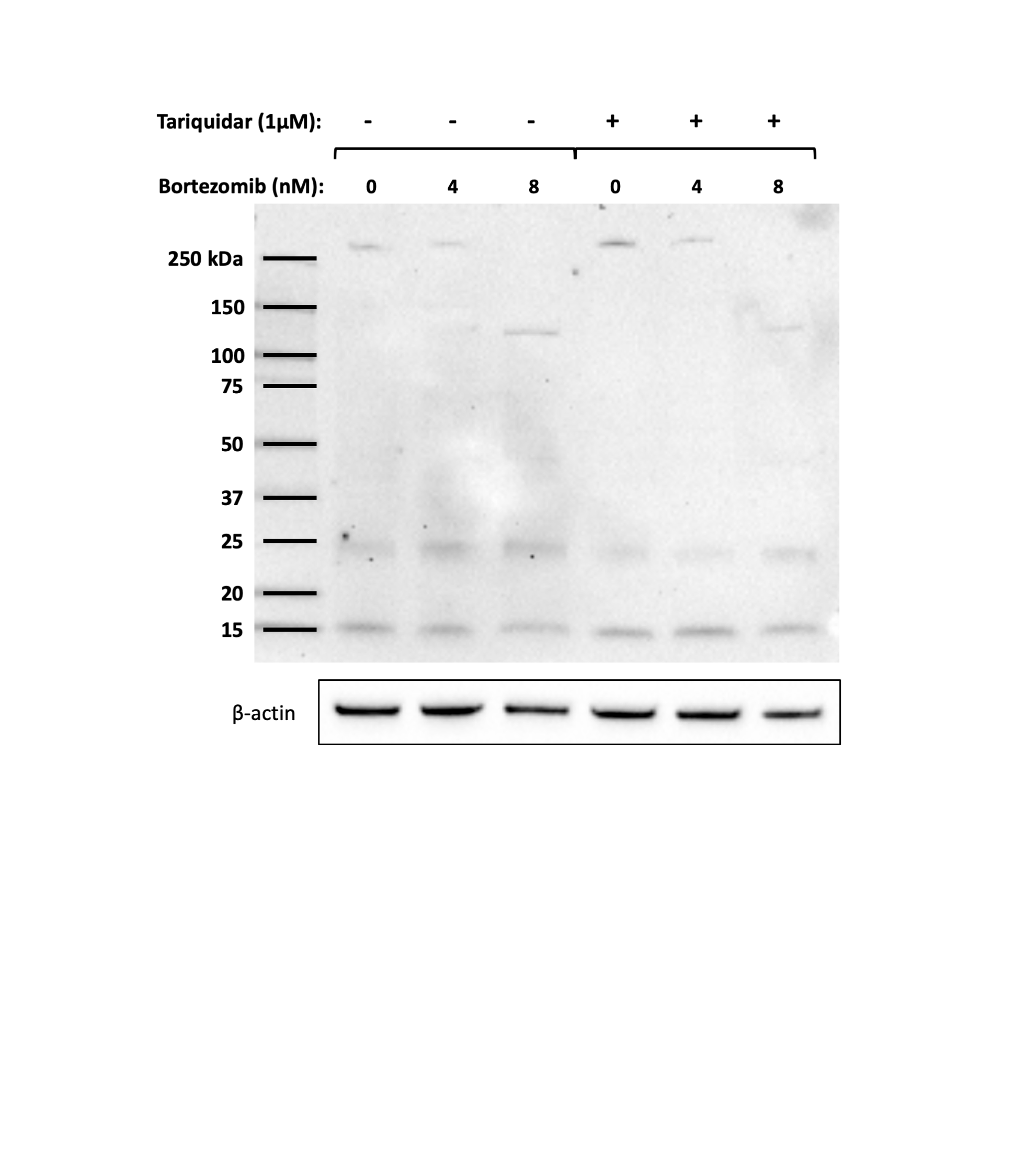
